# Enhancer architecture sensitizes cell specific responses to *Notch* gene dose via a bind and discard mechanism

**DOI:** 10.1101/742908

**Authors:** Yi Kuang, Ohad Golan, Kristina Preusse, Brittany Cain, Joseph Salomone, Ian Campbell, FearGod V. Okwubido-Williams, Matthew R. Hass, Natanel Eafergan, Kenneth H. Moberg, Rhett A. Kovall, Raphael Kopan, David Sprinzak, Brian Gebelein

## Abstract

Notch pathway haploinsufficiency can cause severe developmental syndromes with highly variable penetrance. Currently, we have a limited mechanistic understanding of phenotype variability due to gene dosage. Here, we show that inserting a single enhancer containing pioneer transcription factor sites coupled to Notch dimer sites can unexpectedly induce a subset of *Drosophila Notch* haploinsufficiency phenotypes in an animal with wild type *Notch* gene dose. Mechanistically, this enhancer couples Notch DNA binding to degradation in a Cdk8-dependent, transcription-independent manner. Using mathematical modeling combined with quantitative trait and expression analysis, we show that tissues requiring long duration Notch signals are more sensitive to perturbations in Notch degradation compared to tissues relying upon short duration processes. These findings support a novel “bind and discard” mechanism in which enhancers with specific binding sites promote rapid Notch turnover, reduce Notch-dependent transcription at other loci, and thereby sensitize tissues to gene dose based upon signal duration.

## INTRODUCTION

Haploinsufficiency, or the inability to complete a cellular process with one functional allele of a given gene, manifests in tissue and organ defects with variable penetrance and severity (Wilkie, 1994). For example, *Notch* (*N*) haploinsufficiency, which was discovered in *Drosophila* in 1914, causes a variety of tissue-specific defects that can vary greatly in penetrance and expressivity in flies (Mohr, 1919). Notch pathway haploinsufficiency was subsequently observed in mammals, as *Notch1* heterozygous mice have heart valve and endothelium defects (Nigam and Srivastava, 2009), whereas *Notch2* heterozygotes have defects in bone, kidney and marginal zone B cells (Isidor, et al., 2011; Simpson, et al., 2011; Witt, et al., 2003). A single allele of *NOTCH2* or the *JAG1* ligand can also cause pathological phenotypes in humans, as heterozygosity of either gene can result in a variably penetrant developmental syndrome known as Alagille (McDaniell, et al., 2006; Li, et al., 1997; Oda, et al., 1997). Thus, *Notch* gene dose sensitivity has been observed in a variety of Notch-dependent tissues in both humans and animals. Unfortunately, we currently lack a mechanistic understanding of what causes some tissues to be highly sensitive to *Notch* gene dose and what factors impact the variable penetrance and severity of *Notch* haploinsufficiency phenotypes (Simpson, et al., 2011; McDaniell, et al., 2006; Krebs, et al., 2004; Witt, et al., 2003).

Molecularly, Notch signaling is initiated by ligand-induced proteolysis of the Notch receptor to release the Notch intracellular domain (NICD) from the membrane (Kovall, et al., 2017; Bray, 2016). NICD subsequently transits into the nucleus, binds to the <u>C</u>bf1/<u>S</u>u(H)/<u>L</u>ag1 (CSL) transcription factor (TF) and the adaptor protein Mastermind (Mam), and induces gene expression via two types of DNA binding sites: independent CSL sites that bind monomeric NICD/CSL/Mam (NCM) complexes, and <u>S</u>u(H) <u>p</u>aired <u>s</u>ites (SPS) that are oriented in a head-to-head manner to promote cooperative binding between two NCM complexes (Kovall, et al., 2017; Bray, 2016). Once bound to an enhancer, the NCM complex activates transcription of associated genes via the P300 co-activator. Thus, the production of NICD is converted into changes in gene expression that ultimately regulate cellular processes during development.

Haploinsufficiency of Notch receptor and ligand encoding genes in animals from flies to humans suggests that decreased gene dosage results in a sufficiently large decrease in NICD production to cause phenotypes in a subset of tissues. There is also growing evidence that genetic changes that reduce NICD degradation can alter signal strength with pathological consequences in specific cell types. In the mammalian blood system, for example, Notch1 mutations that remove an NICD degron sequence have been associated with increased NICD levels and the development of T-cell Acute Lymphoblastic Leukemia (T-ALL) in both mice and humans (O’Neil, et al., 2006; Weng, et al., 2004). Intriguingly, NICD turnover via this degron sequence has been directly linked to transcription activation, as the Mam protein interacts with the Cdk8-Mediator submodule, which can phosphorylate NICD to promote its ubiquitylation by the Fbxw7 E3-ligase and degradation by the proteasome (Fryer, et al., 2004; Fryer, et al., 2002). Accordingly, gene mutations that lower Cdk8 activity have also been associated with increased NICD levels and T-ALL initiation and progression (Li, et al., 2014). Thus, perturbations in mechanisms that regulate either NICD production or degradation can induce cell and/or tissue specific phenotypes.

In this study, we use *Drosophila* genetics, quantitative trait and expression analysis, and mathematical modeling to identify two additional factors that impact Notch signal strength and tissue-specific phenotypes. First, we unexpectedly found that an enhancer containing as few as 12 Notch dimer binding sites can induce tissue-specific phenotypes via a Cdk8-Mediator dependent mechanism that can be uncoupled from transcription activation. Second, we built a quantitative signal duration model to reveal how changes in NICD degradation rates preferentially impact long duration Notch-dependent processes, whereas genetic changes in NICD production rates (i.e. *Notch* haploinsufficiency) affect both short and long duration processes. Collectively, these findings provide new insights into how distinct Notch-dependent cellular processes can be differentially impacted by both enhancer architecture and signal duration to induce tissue-specific Notch defects within a complex animal.

## RESULTS

### Enhancers with specific TF binding sites can induce a tissue-specific *Notch* phenotype

To better understand transcriptional responses to *Notch* signals in *Drosophila*, we designed synthetic enhancers with comparable numbers of either CSL monomer (Figure 1A) or SPS dimer sites (Figure 1B, note, 1xSPS equals 2xCSL) (Arnett, et al., 2010; Nam, et al., 2007; Bailey and Posakony, 1995). Since prior studies found that including sites for the Grainyhead (Grh) pioneer TF enhanced Notch reporter activity in *Drosophila* (Furriols and Bray, 2001) and induced chromatin opening (Jacobs, et al., 2018), we generated fly lines containing CSL and SPS reporters with (Figure 1C-E) and without (Figure S1B-C) three copies of a <u>G</u>rh <u>b</u>inding <u>e</u>lement (3xGBE). Surprisingly, flies homozygous for *6xSPS-3xGBE-lacZ* (*6SG-lacZ*) developed the classic *Notch* haploinsufficiency notched-wing phenotype (Figure 1G). In contrast, flies homozygous for *3xGBE* alone (*G-lacZ*), *6xSPS* alone (*6S-lacZ*), or mutated SPSs (*6SmutG-lacZ*) inserted in the same locus were indistinguishable from wild type (Figure S1A-C). To define the *SPS-GBE* binding site features that contribute to wing notching, we tested additional fly lines and found that: i) *6SG-lacZ* caused notched wings when inserted in another locus (Figure S1D-E,H); ii) The relative order of GBE and SPS did not matter (Figure S1E-F,H); iii) The penetrance and severity of wing notching increased as a function of transgene and SPS numbers (Figure 1F-H, and 1J); and iv) Flies with an equal number of Notch monomer (CSL) sites next to 3xGBE did not develop notched wings (Figure 1H-I and 1K). In total, these findings show that adding as few as 12 GBE-associated SPSs into the genome is sufficient to induce a *Notch* haploinsufficiency phenotype in the wing.

**Figure 1.**
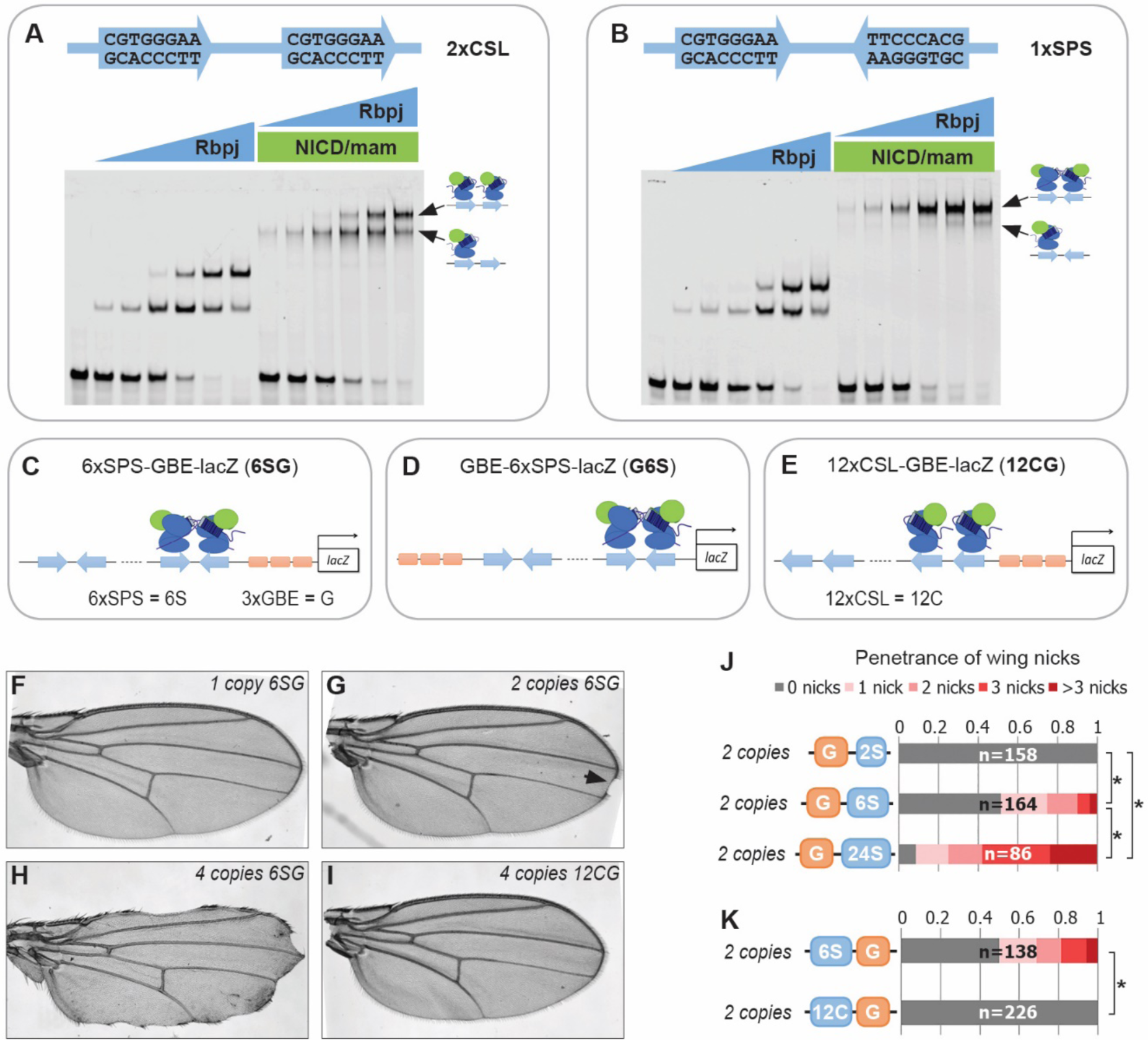
Synthetic Notch enhancers induce a *Drosophila* notched wing phenotype. **A-B**. Electromobility shift assays (EMSAs) using purified NCM proteins and 2xCSL (A) and 1xSPS (B) probes. Arrows highlight bands consistent with one vs two NCM complexes on DNA. **C-E**. Schematics of *6xSPS-3xGBE-lacZ* (*6SG*), *3xGBE-6xSPS-lacZ* (*G6S*) and *12xCSL-3xGBE-lacZ* (*12CG*). **F-I**. Wings from flies with 1 (F), 2 (G, arrowhead denotes a notch) or 4 copies (H) of *6SG-lacZ*, or 4 copies of *12CG-lacZ* (I). **J-K**. Quantified wing notching in flies with indicated transgenes. Proportional odds model tested penetrance and severity differences between *G6S-lacZ* and *G24S-lacZ*. Two-sided Fisher’s exact test assessed penetrance of other genotypes (* p < 0.05). See also Figure S1.

To determine if the *6SG-lacZ* induced wing phenotype could be modified by genetic changes in *Notch* pathway components, we analyzed flies carrying different gene copy numbers of either *N* or the *Hairless* (*H*) co-repressor that antagonizes Notch-mediated gene activation (Morel, et al., 2001; Bang and Posakony, 1992). We found that a single *6SG-lacZ* transgene greatly enhanced the penetrance and severity of wing notching in *N* heterozygotes compared to either genotype alone (Figure 2A-C). Moreover, adding two extra alleles of *N* (4N) or removing one allele of *H* significantly suppressed the notched wing phenotype induced by two copies of *6SG-lacZ* (Figure 2D-H). Thus, wing phenotypes induced by *6SG* can be enhanced or suppressed by changing the gene dose of *N* and *H*, respectively.

**Figure 2.**
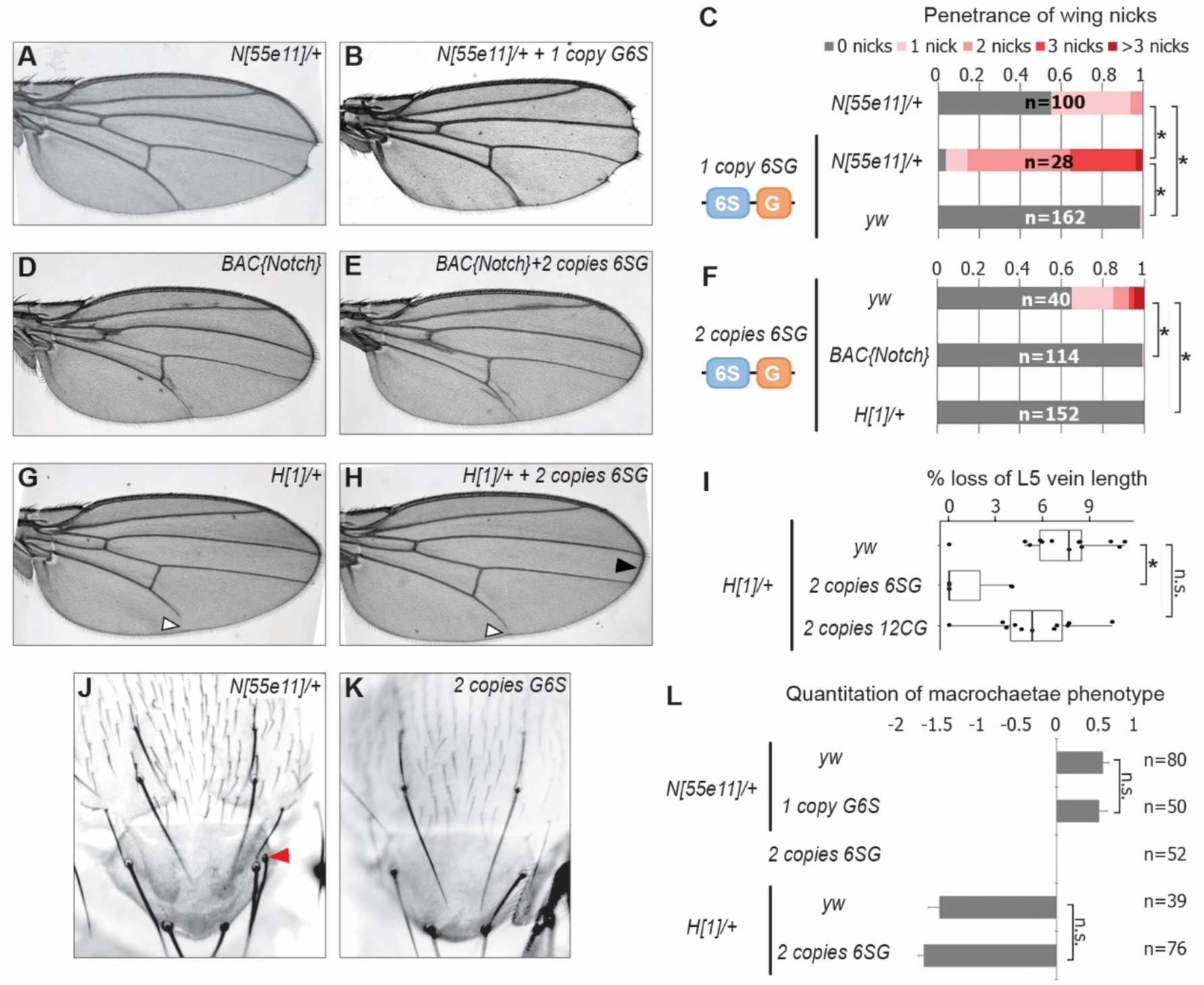
*SPS-GBE* reporters impact wing margin and vein development but not macrochaetae. **A-B**. Wings from *Notch* heterozygotes (*N*^*55e11/+*^) in absence (A) and presence of *G6S-lacZ* (B). **C**. Quantified wing notching in the indicated genotypes. Proportional odds model tested for penetrance/severity differences (* p < 0.05). **D-E**. Wings from flies containing 2 extra *N* alleles (BAC{Notch-GFP-FLAG}) in absence (D) and presence of *6SG-lacZ* (E). **F**. Quantified wing notching in flies with indicated genotypes. Proportional odds model tested *6SG* in *wild-type* and *N*^*55e11/+*^; two-sided Fisher’s exact test compared to *H*^*1/+*^ (* p < 0.05). **G-H**. Wings from *H*^*1/+*^ flies in absence (G) and presence (H) of *6SG-lacZ*. Solid arrowhead highlights loss (G) and rescue (H) of L5 wing vein. Open arrowhead points to rescued *6SG*-induced wing notching phenotype in *H*^*1*/+^ flies. **I**. Quantification of loss of L5 vein in flies with indicated genotypes. Each dot represents a measurement from an individual wing. Two-sided Student’s t-test. In box plots, the line represents median, the box shows interquartile range, and whiskers represent the 1.5 times interquartile range. (* p < 0.05, n.s. not significant) **J-K**. Notum images from a *N*^*55e11/+*^ (J) and *G6S-lacZ* (K) fly. Arrowhead denotes extra macrochaetae in *N*^*55e11/+*^. **L**. Quantification of gained/lost dorsalcentral and scutellar macrochaetae (wild type = 8) in indicated genotypes. Proportional odds model (n.s. not significant).

*N* and *H* haploinsufficiency also cause defects in macrochaetae bristle patterning (Bang, et al., 1991; Shellenbarger and Mohler, 1978) and wing vein development (de Celis and Garcia-Bellido, 1994). Intriguingly, *6SG-lacZ* did not significantly impact macrochaetae formation in either wild type or sensitized *N*^*+/-*^ and *H*^*+/-*^ backgrounds (Figure 2J-L). However, flies carrying two copies of the *6SG-lacZ* Notch-dimer reporter, but not two copies of the *12CG-lacZ* Notch-monomer reporter, significantly suppressed the loss of L5 wing vein observed in *H*^*+/-*^ animals (Figure 2G-I). Altogether, these data demonstrate that coupled *SPS-GBE* sites affect a subset of dose sensitive phenotypes with wing margin cells being the most highly sensitive.

### Cdk8 induces Notch turnover independent from transcription activation

Our findings raise two questions: How does insertion of *SPS-GBE* sites into the genome affect Notch activity, and why only in a subset of Notch-dependent processes? To assess how *SPS-GBE* sites cause notched wings, we screened for mutations that suppressed the wing phenotype and found that removing an allele of each gene of the Cdk8-Mediator submodule (*cdk8* (Loncle, et al., 2007), *cycC* (Loncle, et al., 2007), *kto (Med12)* (Treisman, 2001), or *skd (Med13)* (Treisman, 2001)) or an allele of an E3-ligase that encodes the *Drosophila* homologue of *Fbw7* (*archipelago, ago*) (Moberg, et al., 2001), significantly decreased the penetrance and severity of wing nicking (Figure 3A and Figure S2A). Notably, we observed that in *N* heterozygotes, removing an allele of *cycC, kto, skd* or *ago*, but not *cdk8*, also significantly suppressed wing notching (Figure 3B and Figure S2B). These data are consistent with previous work showing that Cdk8 phosphorylates NICD to promote its ubiquitylation and degradation in mammalian cells (Figure 3C) (Li, et al., 2014; Fryer, et al., 2004). Moreover, the smaller effect of *cdk8* gene dose compared to changes of the other Cdk8-Mediator submodule genes is consistent with studies suggesting *cdk8* is not the limiting factor in the formation of this submodule (Davis, et al., 2013; Knuesel, et al., 2009b). Hence, these findings support the model that lowering the dose of key Cdk8-Mediator submodule genes in *Drosophila* slows NICD turnover and thereby rescues the wing notching phenotype observed in *6SG-lacZ* and *N* heterozygotes.

**Figure 3.**
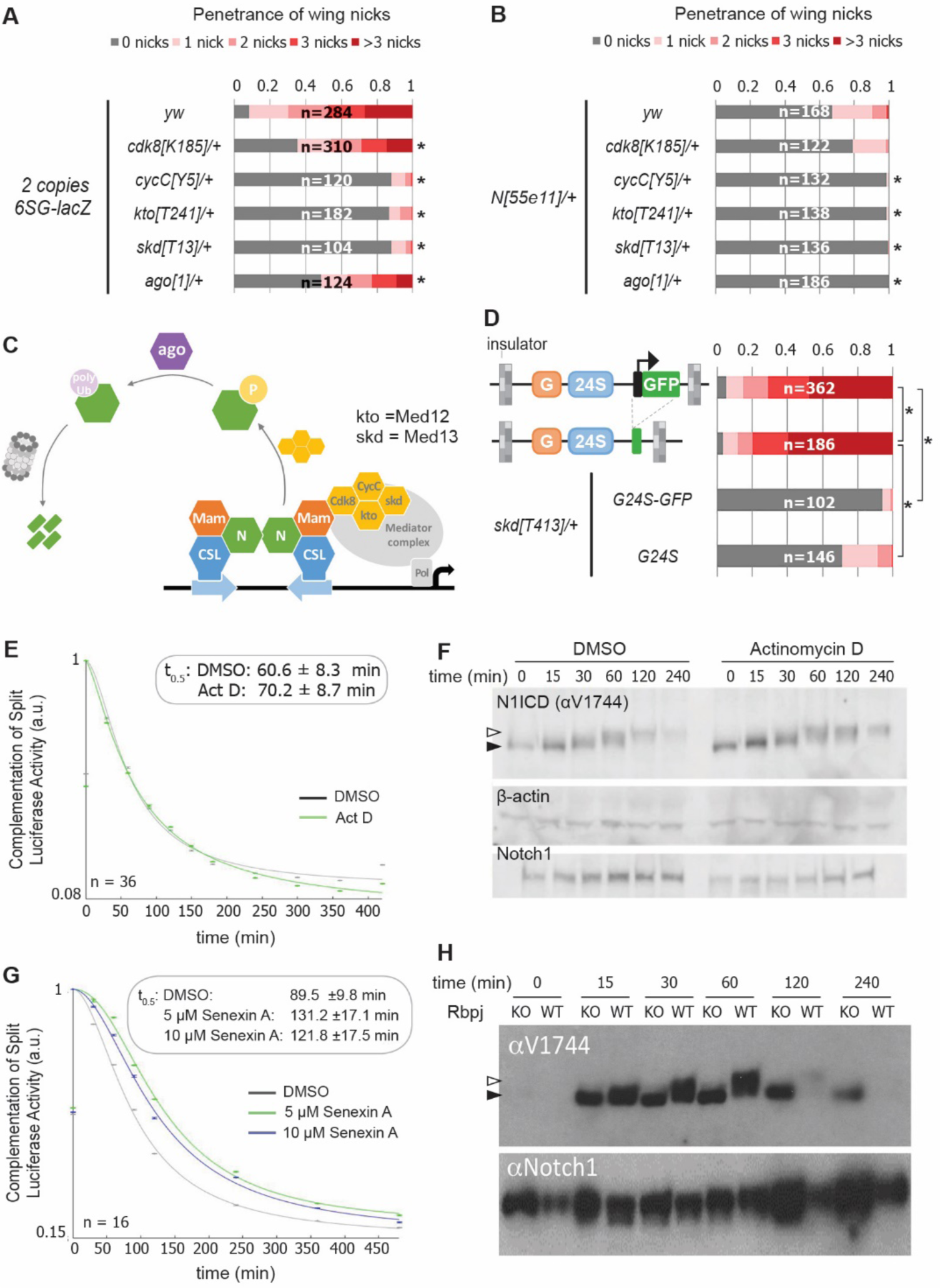
Reducing the activity of the Cdk8-Mediator suppresses the formation of wing notches. **A-B**. Quantified wing notching in *6SG-lacZ* (A) or *N*^*55e11/+*^ (B) in *wild-type* or *cdk8, cycC, kto, skd* or *ago* heterozygotes. Proportional odds model with Bonferroni adjustment tested for significance compared to wild-type (* p < 0.05). **C** Model of Cdk8-mediated NICD degradation. NCM complexes on SPSs recruit Cdk8, CycC, kto and skd. Cdk8 phosphorylates NICD to promote its degradation via Ago and the proteasome (gray cylinder). Cdk8 can also interact with the core Mediator (gray oval). **D** Schematics of promoter-containing and -lacking transgenes at left. Wing notching penetrance and severity at right. Proportional odds model was used to assess significance (* p < 0.05). **E** Rbpj-N1ICD split luciferase assay assessing N1ICD half-life in HEK293T cells treated with DMSO or Actinomycin D. 95% confidence interval noted. **F** Western blot of N1ICD, total Notch1 and β-actin after Notch activation in mK4 cells treated with DMSO or Actinomycin D. **G** Rbpj-N1ICD split luciferase assay assessing N1ICD half-life in HEK293T cells treated with DMSO, 5 μM Senexin A or 10 μM Senexin A. 95% confidence interval noted. **H** Western blot of N1ICD and full-length Notch1 after Notch activation in either *wild-type* (OT13) or *Rbpj*-deficient (OT11) cells. See also Figures S2 and S3.

The Cdk8-Mediator submodule has a complex relationship with promoter transcription (Fant and Taatjes, 2019). Some studies suggest interactions between the Cdk8-Mediator submodule and the core Mediator occludes RNA polymerase recruitment (Knuesel, et al., 2009a) and/or decreases transcription (Pelish, et al., 2015), whereas other studies suggest Cdk8 stimulates transcription (Galbraith, et al., 2013; Donner, et al., 2010). To test the role of the transgene promoter in causing wing nicks, we analyzed flies with promoter-containing (*3xGBE-24xSPS-GFP, G24S-GFP*) or promoter-less transgenes (*3xGBE-24xSPS, G24S*) flanked by insulator sequences (Figure 3D). We found that the wing notching penetrance and severity was similar with both transgenes, and the wing phenotype generated by both was significantly suppressed by removing an allele of *skd* (Figure 3D). These findings suggest that a transcriptionally active promoter is not required to induce wing nicks.

To test the generality of the idea that transcription activation could be uncoupled from NICD degradation, we blocked transcription using actinomycin-D and assessed NICD half-life using a split-luciferase assay in HEK293T mammalian cells (Ilagan, et al., 2011). Importantly, we found that while actinomycin-D effectively inhibited Notch-induced transcription (Figure S3A), it neither altered N1ICD half-life in the split-luciferase assay (Figure 3E), nor altered N1ICD mobility in western blot analysis (Figure 3F). These data suggest that post-translational modification (PTMs) and degradation of N1ICD does not require active transcription. In contrast, inhibiting Cdk8 activity using Senexin-A significantly prolonged N1ICD half-life in this assay (Figure 3G), consistent with Cdk8 playing a key role in regulating NICD turnover. Importantly, we found that N1ICD was stabilized in both mammalian OT11 cells deficient for RBPJ (Figure 3H (Kato, et al., 1997)), and mK4 cells deficient for the three Mastermind-like (MAML) proteins (Figure S3B-C), indicating N1ICD degradation is coupled with NCM complex formation and DNA binding. Moreover, the increased NICD mobility observed in the absence of RBPJ or MAML is consistent with a loss of PTMs (Figure 3H and Figure S3C). Altogether, these data indicate that Cdk8-mediated regulation of NICD degradation requires NCM complex formation on DNA in both mammalian and fly tissues but does not require active transcription.

### Quantitative analysis of Enhancer binding site induced Notch turnover

Our data support a model whereby NCM binding to *SPS-GBE* sites promotes NICD phosphorylation and degradation, and thereby reduces NICD levels in the nucleus (Figure 4A). To obtain a quantitative understanding of how changes in SPS number affects Notch signal strength, we used mathematical modeling and quantitative expression analysis. The model includes a set of biochemical reactions that describe NICD dynamics in the nucleus (bottom, Figure 4A). We initially assume unphosphorylated, unbound NICD (*NICD*_*up,ub*_) enters the nucleus at a constant production rate (*P*_*NICD*_), where it forms NCM complexes that bind DNA. Bound, unphosphorylated NICD (*NICD*_*up,b*_) can be phosphorylated by Cdk8 (*NICD*_*p,b*_) at a rate *k*_*p*_. Similar to the unphosphorylated state, phosphorylated NICD can cycle between *NICD*_*p,b*_ and *NICD*_*p,ub*_. Finally, it is assumed that the degradation rate of *NICD*_*p,ub*_, denoted by Γ_*p*_, is much faster than the degradation rate of *NICD*_*up,ub*_, denoted by Γ_*up*_.

**Figure 4.**
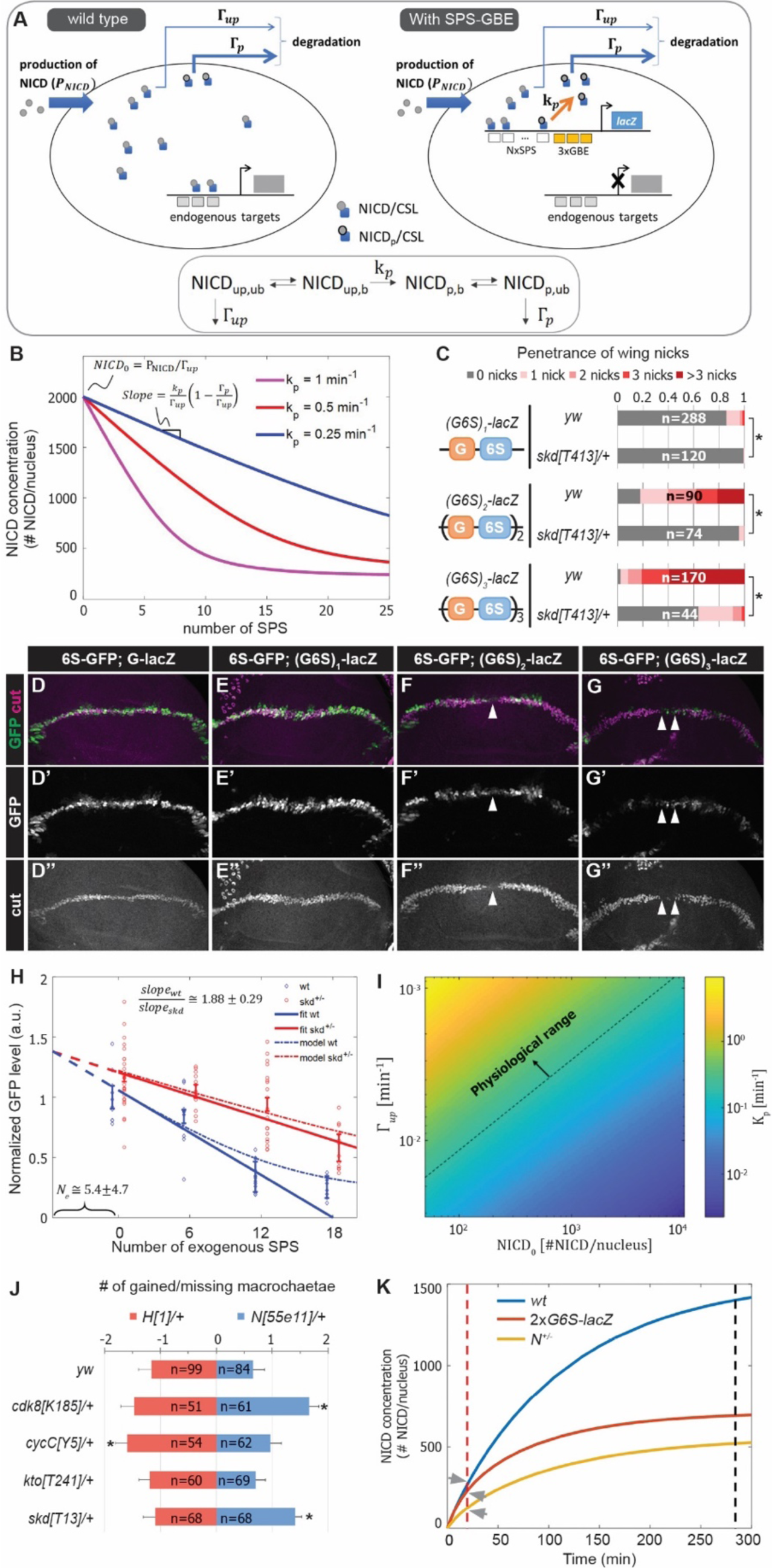
A mathematical model coupling NICD degradation to DNA binding predicts Notch activity and tissue sensitivity. **A** Schematic and equation describing *SPS-GBE* induced turnover of NICD. In both wild type (left) and nuclei with SPS-GBE sites inserted (right), NICD is produced and enters the nucleus at a constant rate (*P*_*NICD*_). NCM complexes form on SPS, where NICD is phosphorylated by Cdk8 at a rate *k*_*p*_. Phosphorylated NICD degrades faster (Γ_*p*_) than unphosphorylated NICD (Γ_*up*_). Subscripts *p, up, b, ub* denote phosphorylated, unphsophorylated, bound, and unbound NICD. **B** Simulations of NICD levels as a function of SPS number. The three curves correspond to simulations with indicated values of *k*_*p*_. NICD starts from a common level (*NICD*_0_) and initially decreases linearly with number of SPS, with a slope proportional to *k*_*p*_. **C** Wing notching penetrance and severity in flies with indicated genotypes. Proportional odds model was used to assess significance (* p < 0.05). **D-G”**. Wing discs from flies containing *6xSPS-GFP* (*6S-GFP*) and either *GBE-lacZ* (*G-lacZ*) or *(G6S)*_*1,2 or 3-*_*lacZ* stained with cut (magenta). **H** Quantified GFP levels in wing discs with increasing SPSs (0, 6, 12, 18 correspond to *(G6S)*_*1,2*_ *or 3-lacZ*) in either wild type (blue) or *skd* heterozygotes (red). Each dot represents average GFP level in margin cells from a single wing disc. Error bars show means and S.E.M for each disc. Solid lines represent linear fit to mean GFP values of first three points of *wild-type* (blue) and four points of *skd* heterozygotes (red). Ratio of slopes is indicated. Effective number of endogenous SPS, *N*_*e*_, is estimated by extrapolating the y axis intersect of dashed lines. **I** Estimated phosphorylation rates by a single SPS, *k*_*p*_. Values of *k*_*p*_ (color-bar) were estimated for a range of values of *NICD*_0_ and Γ_*up*_. Dashed line represents lower limit of the physiological range of kinase activities. **J** Quantified number of gained/lost macrochaetae from indicated genotypes in *N*^*55e11/+*^ (blue bars) or *H*^*1/+*^ (red bars) background. Proportional odds model with Bonferroni adjustment. Data are mean ± 95% confidence interval (* p < 0.05). **K** NICD level simulations as a function of time after Notch activation (at t=0 min) in *wild-type* (blue), *N* heterozygotes (yellow) and *SPS-GBE* flies (red). In tissues with long duration Notch activation (black dash line), *N* heterozygotes and *SPS-GBE* sites similarly reduce NICD levels. In tissues with short duration Notch activation (red dash line), NICD levels are weakly affected by SPS-GBE compared to *N* heterozygotes (arrows). See also Figures S4 and S5.

Analysis of the differential equations corresponding to these reactions generated several predictions. First, our model predicts steady state NICD levels will initially decrease linearly as the number of SPSs increases and then saturate for high numbers of SPSs (Figure 4B). Importantly, the linear regime of the slope describing NICD degradation is expected to be proportional to the Cdk8 phosphorylation rate, *k*_*p*_. Accordingly, if there is no dosage compensation in Cdk8-Mediator submodule heterozygotes, the model’s second prediction is that the slope of the wild-type curve should be twice that of the heterozygous mutant curve.

To test these predictions, we measured Notch signal strength in wing margin cells in the presence and absence of *SPS-GBE* transgenes. To systematically vary SPS numbers, we created fly lines containing one, two or three (*3xGBE-6xSPS*) cassettes in front of a single *lacZ* gene (*(G6S)n-lacZ*). Analysis of flies carrying a single copy of the *(G6S)n-lacZ* transgenes revealed enhanced penetrance and severity of wing notching as the number of *G6S* cassettes increased, and all were significantly suppressed by removing one *skd* allele (Figure 4C). Because direct measurement of nuclear NICD levels *in vivo* is very challenging (Couturier, et al., 2012), we monitored NICD levels in wing margin cells indirectly via GFP expression from an independent *6xSPS-GFP* (*6S-GFP*) reporter that is highly sensitive to changes in *Notch* gene dose (Figure S4). We found that GFP levels decreased as a function of added *GBE-SPS* sites (Figure 4D-H). Simultaneous analysis of Cut, an endogenous *Notch* target required for maintaining wing margin fate (Micchelli, et al., 1997; Neumann and Cohen, 1996), revealed a loss of wing margin fate in a subset of *(G6S)*_*2*_*-lacZ* and *(G6S)*_*3*_*-lacZ* cells (arrowheads in Figure 4F-G), consistent with the notched wing phenotype observed in these animals.

Analysis of *6S-GFP* expression revealed an approximately linear decrease in GFP as the number of *G6S* cassettes is increased (Figure 4H, blue markers). Moreover, removing an allele of *skd* significantly increased *6S-GFP* expression, resulting in a shallower slope relative to wild type flies with the same *(G6S)-lacZ* transgene (Figure 4H, red markers). The ratio between slopes as calculated by linear regression analysis of GFP levels in wild type and *skd* heterozygotes (solid lines in Figure 4H) was 1.88±0.29, in agreement with the predicted 2-fold change in the absence of dosage compensation. Interestingly, the two curves did not intersect at the y-axis, reflecting a cumulative reduction in NICD phosphorylation and degradation rates at endogenous sites; an interpretation supported by the observation that Cdk8-Mediator submodule heterozygotes ameliorate *Notch* heterozygote induced wing notching phenotypes (Figure 3B). We used this observation to estimate the magnitude of the cumulative genomic effect by extrapolating the crossing point of the two curves (dashed lines in Figure 4H). The lines crossed at negative *N*_*e*_ = 5.4 ± 4.7 *SPS-GBE* sites. This value means that the cumulative effect on NICD stability of all sites in the genome (*N*_*e*_) is equal to that of ∼5 highly active synthetic *SPS-GBE* sites.

Next, we used the model to calculate Cdk8 phosphorylation rates, *k*_*p*_, needed at *SPS-GBE* sites to lower NICD concentrations and induce wing notching phenotypes. In the linear regime, *k*_*p*_ (in 1/min units) can be calculated from the measured slope, *slope*_*wt*_, for different values of *NICD*_0_, Γ_*up*_, and Γ_*p*_ (see Figure 4B and Experimental Procedures for derivation). We used a plausible range of NICD concentrations, *NICD*_0_, (between 10^2^-10^4^ molecules per nucleus) and degradation rates Γ_*up*_ (between 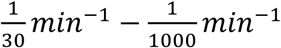, see methods for parameter estimation). This analysis provided a range for the Cdk8 catalytic activity on *SPS-GBE* sites between 2 × 10^−3^ < *k*_*p*_ < 10 *min*^−1^ (Figure 4I). Because kinase activity rates typically range between 10^−1^to 10^3^ *min*^−1^ (Davidi, et al., 2016; Koivomagi, et al., 2011; Good, et al., 2009), a large portion of the parameter space falls within the physiological regime of known kinase activities (above the dashed line in Figure 4I). In fact, the calculated *k*_*p*_ values are at the low end of the physiological range (of the order of ∼1/min), suggesting that even modest phosphorylation rates could produce the observed reduction in Notch signal strength, and ultimately, wing notching phenotypes induced by *SPS-GBE* sites.

### Altered NICD degradation sensitizes tissues requiring long duration signals

Molecularly, *Notch* haploinsufficiency is due to decreased NICD production, whereas the phenotype caused by *SPS-GBE* sites is due to enhanced NICD turnover. This difference in mechanism might explain why *SPS-GBE* sites fail to impact another known *N* dose sensitive tissue, the macrochaetae sensory bristles (Figure 2K-L). These data support the idea that wing margin cells are sensitive to changes in both NICD production and degradation, whereas macrochaetae formation is preferentially sensitive to changes in NICD production. To further test this hypothesis, we analyzed macrochaetae formation in compound heterozygotes for Cdk8-Mediator submodule genes and *Notch*. In contrast to the observed suppression of wing notching (Figure 3B), we found that removing an allele of each Cdk8-Mediator submodule gene did not significantly suppress the *Notch* haploinsufficiency of extra macrochaetae (Figure 4J). In fact, macrochaetae numbers further increased in *N;cdk8* and *N;skd* compound heterozygotes, which is opposite of the predicted outcome if slowing NICD degradation significantly elevated NICD levels during macrochaetae formation (Figure 4J). As a second test to determine if changes in the Cdk8-Mediator submodule could alter macrochaetae, we also analyzed *H* heterozygotes that generate too few macrochaetae due to increased Notch activity (Morel, et al., 2001; Maier, et al., 1997). In this genetic background, removing an allele of either *skd* or *cdk8* did not significantly alter macrochaetae formation (Figure 4J). Thus, these data suggest that macrochaetae formation is relatively insensitive to changes in NICD degradation rates.

A potential explanation for the tissue-specific response to *SPS-GBE* sites and Cdk8 Mediator submodule heterozygotes could be the distinct temporal requirements for Notch activation in each tissue. Maintenance of wing margin identity is a continuous process at least 48 hours long (de Celis, et al., 1996; Shellenbarger and Mohler, 1978), whereas macrochaetae formation requires Notch input over a short time period (< 30 min) (Barad, et al., 2010). To explore the relationship between Notch signal duration and sensitivity to changes in NICD production/degradation rates, we modeled the dynamics of NICD accumulation as a function of time in wild-type, *N* heterozygotes, and flies homozygous for *G6S-lacZ* (Figure 4K). We assume that *N* heterozygotes lower NICD production (*P*_*NICD*_) by one half without impacting NICD degradation, whereas *SPS-GBE* sites do not affect *P*_*NICD*_ but increase NICD degradation as a function of SPS number. In a scenario where nuclear NICD reaches steady state (Figure 4K, black dash line), both the *SPS-GBE* loci and *N* heterozygotes significantly decrease NICD levels. In contrast, if Notch signals are only required for a short time period, changes in degradation rates do not significantly alter NICD levels relative to the impact of losing a *N* allele (Figure 4K, arrows). We note that this conclusion is robust over a broad range of potential NICD production and degradation rates (Figure S5). Moreover, the model is consistent with the results observed using genetic changes in Cdk8-Mediator submodule gene dose – altering NICD degradation selectively impacts long duration events (wing margin) and not short duration events (lateral inhibition during macrochaetae specification).

## DISCUSSION

Our results show that simply increasing the number of clustered Notch dimer sites (SPS) linked to sites for the Grh pioneer TF (GBE) can cause a tissue-specific *N* haploinsufficiency phenotype via a Cdk8-dependent mechanism. These findings have several important implications for both enhancer biology and the mechanisms regulating Notch signal strength in specific tissues. First, the proposed mechanism links degradation of the Notch signal (NICD) with its binding to specific loci (accessible SPSs) in a manner that can be uncoupled from transcription activation. This “bind and discard” mechanism reveals an unexpected global link between all exposed binding sites in the epigenome, such that the collective “drain” loci can reduce Notch-dependent transcription at other loci in the same nucleus. In fact, inserting a single enhancer containing as few as 12 Notch dimer sites was sufficient to decrease signal strength in wing margin cells to a level similar as removing an entire *Notch* allele.

Second, our data supports the idea that not all Notch binding sites are equally capable of marking NICD for degradation, and that enhancer architecture plays a key role in modulating NICD turnover. For instance, only Notch dimer but not Notch monomer sites are sufficient to generate phenotypes, and even Notch dimer enhancers differ in their ability to induce phenotypes based on the absence/presence of Grh sites. Since Grh binding is sufficient to increase chromatin opening (Jacobs, et al., 2018), these findings suggest enhancer accessibility alters the rate at which NICD is metabolized by Notch dimer sites. Intriguingly, published ChIP data (Nevil, et al., 2017) reveals Grh binds extensively to the *Enhancer of Split* (*E(spl)*) locus that has numerous SPS-containing *Notch* regulated enhancers (Cave, et al., 2011; Cooper, et al., 2000; Nellesen, et al., 1999). Thus, these data highlight a potential mechanism by which enhancer architecture (i.e. Notch dimer vs monomer sites) and epigenetic “context” (i.e. accessibility due to pioneer TF binding) can fine tune the global Notch response in different tissues.

Third, we propose that the differential sensitivity of Notch-dependent tissues to changes in NICD degradation rate (i.e. *SPS-GBE* sites or Cdk8-Mediator heterozygotes) or NICD production rates (*N* heterozygotes) reflects the temporal requirement for Notch signal duration. Intuitively, these findings suggest a mechanism underlying cell-specific context that may have implications for both developmental processes and tumorigenesis. For example, mutations in the NICD PEST domain that decouple DNA binding and degradation are common in T-cell acute lymphoblastic leukemia (T-ALL) (Weng, et al., 2004), and CycC (CCNC) functions as a haploinsufficient tumor suppressor gene in T-ALL, at least in part, by stabilizing NICD (Li, et al., 2014). Indeed, T-ALL cells are “addicted” to Notch and are thus dependent on a long duration signal (Severson, et al., 2017). As ∼30% of Notch target genes in T-ALL use SPS containing enhancers, our findings provide insight into how Notch PEST truncations and CCNC heterozygotes would each promote tumorigenesis by slowing Cdk8-mediated NICD turnover on SPS enhancers. It is also important to note that the Cdk8-Mediator submodule interacts with many genomic loci (Pelish, et al., 2015), suggesting this mechanism may be quite general and apply to transcription factors beyond Notch. Future studies focused on enhancers that recruit the Cdk8-Mediator submodule and other Notch-dependent cellular processes will help reveal how the temporal requirements for nuclear activities contributes to both normal development and disease states.

## ACKNOWLEDGEMENTS

We thank members of the Gebelein, Sprinzak and Kopan labs for comments on this work. We thank Drs. Stephen Blacklow, Masato Nakafuku, Kenneth Campbell, and Ertugrul Ozbudak for comments on the manuscript and/or project. The cdk8[K185] and cycC[Y5] alleles were gifts from Dr. Jessica Treisman (New York University). We thank the *Drosophila* stock centers and Developmental Studies Hybridoma Bank (DSHB) for fly stocks and antibody reagents. This work was supported by NSF/BSF grant #1715822 (B.G., D.S., and R.A.K.), NIH grant CA163653 (R.K.) and NIH grant CA178974 (R.A.K.).

## AUTHOR CONTRIBUTIONS

This scientific study was conceived and planned by Y.K., O.G., R.K., D.S., and B.G. The transgenic constructs and *Drosophila* reagents were generated by Y.K., B.C., K.H.M., and B.G. *Drosophila* experiments were performed by Y.K., B.C., and F.O.W. Mammalian cell culture experiments were performed by K.P. and M.H. Protein purification and gel shift analysis was performed by I.C., J.S., and R.A.K. Data analysis was performed by Y.K., O.G., K.P., N.E., R.K., D.S., and B.G. Mathematical modeling was performed by O.G. and D.S. The manuscript was written by Y.K., R.K., D.S., and B.G.

## EXPERIMENTAL PROCEDURES

### Reporter design, molecular cloning, and transgenic fly generation

All synthetic Notch enhancer sequences contain high affinity Su(H) binding sites (CGTGGGAA). Notch monomer sites (CSL) were placed 17bps apart in a head-to-tail manner to permit independent binding of NCM complexes. Notch dimer sites (SPS) were spaced 15bps apart in a head-to-head orientation to enable cooperative dimerization between adjacent NCM complexes. Intervening sequences were designed to exclude known binding sites for other *Drosophila* TFs using the *cisBP* website (Weirauch, et al., 2014). The 6xSPSmut sequence is identical to 6xSPS except for two nucleotide changes (CG<u>A</u>GG<u>C</u>AA) in each binding site. The 2xSPS, 6xSPS, 6xSPSmut and 12Xcsl sequences were synthesized by GenScript as either complimentary oligonucleotides (2xSPS) or double stranded DNA (6xSPS, 6xSPSmut and 12xCSL; complete sequences are listed below). Cloning was facilitated by including flanking EcoR1 and BglII sequences. Annealed oligonucleotides or double stranded DNA fragments were cloned into either *placZ-attB* or *3xGBE-placZ-attB* (Uhl, et al., 2016) and sequence confirmed. To concatenate 6xSPS into larger arrays, a shuttle vector was used to generate a BamH1-6xSPS-BglII-Not1 fragment for reiterative cloning into vectors digested with BglII/NotI (BglII/NotI permits cloning a BamH1/NotI fragment, which can be repeated as desired). To create the *3xGBE-24xSPS-lacZ* vector, a *24xSPS* fragment (generated in the shuttle vector) was cloned into the *3xGBE-lacZ* vector. To make the promoter containing *3xGBE-24xSPS*-*GFP* vector, we inserted *3xGBE-24xSPS* into *pHStinger-attB* (Barolo, et al., 2000). To generate a promoterless *3xGBE-24xSPS* construct, the promoter and GFP encoding sequences were removed from *3xGBE-24xSPS-GFP* by KpnI/SpeI digest, blunted with T4 DNA polymerase, and ligated. To generate the *(3xGBE-6xSPS)n-lacZ* (n=2,3) constructs, the *3xGBE-6xSPS* fragment was iteratively cloned into the *3xGBE-6xSPS-lacZ* plasmid. The *6xSPS-GFP* reporter used to measure Notch transcription responses was generated by cloning 6xSPS into the *pHStinger-attB* vector. All transgenic fly lines were generated by phiC31 recombinase integration into 22A, 51C or 86Fb loci of the *Drosophila* genome as indicated (Rainbow Transgenic Flies, Inc).

### Synthetic enhancer DNA and probe design

Enhancer sequences used in the transgenic reporter vectors and the DNA probes used in electromobility shift assays (EMSAs) are listed in FASTA format. Restriction enzyme sites (RE) and/or RE overhangs used for cloning are highlighted in yellow, the Su(H) binding sites are highlighted in cyan, and point mutations are bold and underlined.

>6xSPS

GAATTCAGCTACGTGGGAAAGGAGCAAACTGCGTTTCCCACGTTCGCAGGGCAGCTACGTGGGAAAGGAGCAAACTGCGTTTCCCACGTTCGCAGGGCAGCTACGTGGGAAAGGAGCAAACTGCGTTTCCCACGTTCGCAGGGCAGCTACGTGGGAAAGGAGCAAACTGCGTTTCCCACGTTCGCAGGGCAGCTACGTGGGAAAGGAGCAAACTGCGTTTCCCACGTTCGCAGGGCAGCTACGTGGGAAAGGAGCAAACTGCGTTTCCCACGTTCGCAGGGCAGATCT

>6xSPS-mut

GAATTCAGCTACG<u>**A**</u>GG<u>**C**</u>AAAGGAGCAAACTGCGTTT<u>**G**</u>CC<u>**T**</u>CGTTCGCAGGGCAGCT ACG<u>**A**</u>GG<u>**C**</u>AAAGGAGCAAACTGCGTTT<u>**G**</u>CC<u>**T**</u>CGTTCGCAGGGCAGCTACG<u>**A**</u>GG<u>**C**</u>AA AGGAGCAAACTGCGTTT<u>**G**</u>CC<u>**T**</u>CGTTCGCAGGGCAGCTACG<u>**A**</u>GG<u>**C**</u>AAAGGAGCAAAC TGCGTTT<u>**G**</u>CC<u>**T**</u>CGTTCGCAGGGCAGCTACG<u>**A**</u>GG<u>**C**</u>AAAGGAGCAAACTGCGTTT<u>**G**</u>CC <u>**T**</u>CGTTCGCAGGGCAGCTACG<u>**A**</u>GG<u>**C**</u>AAAGGAGCAAACTGCGTTT<u>**G**</u>CC<u>**T**</u>CGTTCGCAG GGCAGATCT

>12xCSL

GAATTCGCCCTGCGAACGTGGGAAACCTAGGCTAGAGGCACCGTGGGAAACTGCCTGCCCTGCGAACGTGGGAAACCTAGGCTAGAGGCACCGTGGGAAACTGCCTGCCCTGCGAACGTGGGAAACCTAGGCTAGAGGCACCGTGGGAAACTGCCTGCCCTGCGAACGTGGGAAACCTAGGCTAGAGGCACCGTGGGAAACTGCCTGCCCTGCGAACGTGGGAAACCTAGGCTAGAGGCACCGTGGGAAACTGCCTGCCCTGCGAACGTGGGAAACCTAGGCTAGAGGCACCGTGGGAAACTGCCTAGATCT

>2xSPS_Cloning_oligonucletide#1

aattcAGCTACGTGGGAAAGGAGCAAACTGCGTTTCCCACGTTCGCAGGGCAGCTACGTGGGAAAGGAGCAAACTGCGTTTCCCACGTTCGCAGGGCa

>2xSPS_Cloning_oligonucleotide#2

gatctGCCCTGCGAACGTGGGAAACGCAGTTTGCTCCTTTCCCACGTAGCTGCCCTGCGAACGTGGGAAACGCAGTTTGCTCCTTTCCCACGTAGCTg

>1xSPS_EMSA_oligonucleotide

GCTACGTGGGAAAGGAGCAAACTGCGTTTCCCACGTTCGTAGTGCGGGCGTGGCT

>2xCSL_EMSA_oligonucleotide

CGAACGTGGGAAACCTAGGCTAGAGGCACCGTGGGAAACTAGTGCGGGCGTGGCT

>EMSA_5’IRDye-700_complementary_oligonucleotide

AGCCACGCCCGCACT

### Fly husbandry

The following alleles were obtained from the Bloomington *Drosophila* Stock Center: PBac{N-GFP.FLAG}VK00033 (stock #38665), N[55e11] (#28813), H[1] (#515), kto[T241] (#63126), kto[T631] (#63125), skd[T13] (#63123) and skd[T413] (#63124). cdk8[K185] and cycC[Y5] alleles were gifts from Professor Jessica Treisman (Treisman, 2001). ago[1] and ago[3] alleles were previously described (Moberg, et al., 2001). Flies were maintained under standard conditions with all genetic crosses, phenotyping and gene expression assays performed at 25°C. The detailed genetic crosses needed to generate the progeny in each Figure are listed in Table S1.

### Genetic assays

To analyze the wing notching and macrochaetae phenotypes, flies of the appropriate genotypes were mated in cornmeal-containing vials and transferred to fresh food every day to avoid overcrowding. Offspring of the listed genotypes were selected and the number of nicks on each wing was recorded and/or the number of dorsocentral and scutellar macrochaetae was counted. A Fisher’s exact test was used to determine significance between samples when penetrance was being assessed (i.e. no phenotype versus a phenotype), whereas a proportional odds model was used to determine significance when analysis included phenotype severity. For L5 wing vein length, fly wings of the proper genotypes were dissected, mounted on glass slides and imaged using a Nikon NiE upright widefield microscope. The total length of the presumptive L5 vein and the vein-missing gap were measured using Imaris software. A student’s t-test was used to determine significance.

### GFP reporter assays in larval imaginal wing discs

To systematically assess Notch transcription responses in larval wing imaginal discs, animals homozygous for *6xSPS-GFP*_*22A*_ and either *3xGBE-lacZ, 3xGBE-6xSPS-lacZ, (3xGBE-6xSPS)*_*2*_*-lacZ*, or *(3xGBE-6xSPS)*_*3*_*-lacZ* were mated to either *yw* (wild type) or *skd[T413]/TM6B* males. Imaginal discs from male non-TM6B wandering 3^rd^ instar larvae (*skd[T413]* heterozygotes) were dissected and fixed in 4% formaldehyde for 15 min. Samples were subsequently washed 4 times with PBX (0.3% Triton X-100 in PBS) and incubated with an antibody that recognizes the Cut antigen (mouse 1:50, DSHB) followed by a fluorescent-conjugated secondary antibody (Alexa Fluor, Molecular Probes). For quantitative purposes, at least 8 imaginal discs were analyzed for each genetic condition tested and the entire wild type imaginal disc series was harvested, fixed and imaged at the same time using a Nikon A1R inverted confocal microscope (40x objective). For the *skd* heterozygote series, a set of wild type imaginal discs with *3xGBE-lacZ* was performed simultaneously to normalize the responses between series. All imaging was performed with constant settings for GFP levels, and GFP pixel intensity in wing margin cells was determined from Z-stack images using Imaris software. A two-sided Student’s t-test was used to determine significance between samples.

### Protein purification and electrophoretic mobility shift assay (EMSA)

For recombinant protein purification, constructs that correspond to mouse RBPJ (aa 53-474), mouse N1ICD (aa 1744-2531), and human SMT3-MAML1 (aa 1-280) were expressed and purified from bacteria using a combination of affinity (Ni-NTA or Glutathione), ion exchange, and size exclusion chromatography as previously described (Friedmann, et al., 2008). Purified proteins were confirmed by SDS-PAGE with Coomassie blue staining and concentrations were measured by absorbance at UV280 with calculated extinction coefficients. Electromobility shift assays (EMSAs) were performed as previously described using native polyacrylamide gel electrophoresis (Uhl, et al., 2016; Uhl, et al., 2010). Fluorescent labeled probes (3.4nM) were mixed with increasing concentrations of purified CSL (RBPJ) protein in 2-fold steps (from 9.4 to 300nM) with or without purified NICD and Mastermind (Mam) proteins at 550nM. Acrylamide gels were imaged using the LICOR Odyssey CLx scanner.

### Split luciferase assay and half-life estimations

Stability of mammalian N1ICD was analyzed using a previously described Luciferase complementation assay (Ilagan and Kopan, 2014; Ilagan, et al., 2011). HEK293T cells with stable expression of CLuc-RBPjK and NOTCH1-NLuc were cultured for 8 hours with 50 nM Actinomycin D to block transcription or 4 hours in the presence of the inhibitor Senexin A to block CDK8-mediated phosphorylation. Cells were activated for 10 min with 0.05% Trypsin-EDTA or with Trypsin only as a negative control and transferred to Poly-D-Lysine coated black 96-well plates with 40,000 cells/well. Cells were cultured at 37°C for 1 hour to let cells attach. Medium was changed to Opti-MEM (Gibco) with 150 μg/ml D-Luciferin (Goldbio), the substrate was added fresh before each measurement. Luciferase signals were measured using the IVIS Lumina LT system and were normalized to Trypsin treated controls and each well separately to the signal counts at t=0. To confirm activity of Actinomycin D, mK4 cells with a stably integrated a 6xSPS-NanoLuc reporter or a 0xSPS-NanoLuc construct as control were used. Cells were cultured for 8 hours with 50 nM Actinomycin D or 0.1% DMSO and activated for 10 min with 0.05% Trypsin-EDTA (Gibco). Nano-Luc activity was measured after 3 hours using the Nano-Glo® Luciferase Assay (Promega) and imaged with the IVIS Lumina LT system. Since the decay curves in Figure 3G did not exhibit simple exponential decay, we calculated the half-life values by fitting the luciferase activity values to a decreasing Hill function, 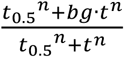 where t is time, *t*_0.5_ is the half life, *bg* is the background level, and *n* is the Hill coefficient. The fitting was performed using least mean square fitting algorithm in Matlab. The error on half-life is the 95% confidence interval.

### Western blot analysis

For Western blot analysis of mammalian N1ICD, mK4 cells were cultured 8 hours with 50 nM Actinomycin D (Sigma-Aldrich, A1410) or 0.1% DMSO in medium before activation of NOTCH with 0.05% Trypsin-EDTA (Gibco) for 10 min. Cells were harvested at different timepoints, lysed in RIPA and sonicated. Equal amounts were loaded on 6% Acrylamide gels for SDS-PAGE and blotted on Nitrocellulose membranes (GE healthcare). Membranes were blocked in 5% dry milk powder in PBS with 0.1% Tween and incubated with anti-N1ICD (Val1744, CST, 1:1000), anti-Notch1 (D1E11, CST, 1:1000) and anti-β-actin (Sigma-Aldrich, 1:4000) overnight at 4°C. After incubation with HRP-conjugated secondary antibodies (GE Healthcare, 1:5000), signals were detected using SuperSignal™ Femto West Chemoluminescent Substrate (Thermo Fisher Scientific) and imaged with the BioRad ChemiDoc system.

Mouse embryonic fibroblast cells deficient for RBP-JK (OT-11, (Kato, et al., 1997)) or wild-type control (OT-13) cells were collected at various times after trypsin/EDTA treatment from confluent wells of a 12 well plate. The cells were washed with PBS, lysed in 100 ul of RIPA + protease inhibitors and 100 ul of 2X sample buffer, and DNA was sheared with a needle. Equivalent amounts of lysate were run on an SDS polyacrylamide gel, transferred to nitrocellulose, blotted for active (Val1744, CST, 1:1000) or total Notch1 protein (D1E11, CST, 1:1000), and developed as described above.

We generated mK4 cells deficient for all mastermind-like proteins through CRISPR Cas9 mediated deletion of exon 1 of Mastermind-like 1, 2, and 3. Briefly, parental mK4 cells were simultaneously transfected with PX458 and PX459 containing guide RNAs flanking exon1 for each of the Mastermind-like genes and subjected to selection with puromycin for 2 days. Surviving clones were picked the following week using cloning disks and screened by Western blot to identify clones that lacked expression of Mastermind-like 1, 2, and 3.

### Mathematical model

The mathematical model describes the concentrations of the possible states of NICD. These include unphosphorylated unbound NICD, *NICD*_*up,ub*_, unphosphorylated bound, *NICD*_*up,ub*_, phosphorylated bound, *NICD*_*p,b*_, and phosphorylated unbound, *NICD*_*p,ub*_. The dynamic equations described by the set of biochemical reactions presented in Figure 4a are:

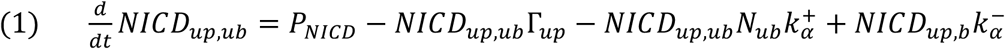

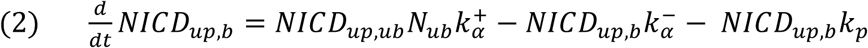

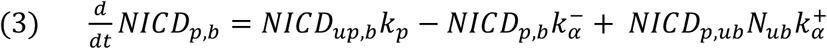

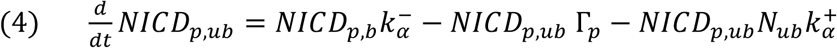

Here, *P*_*NICD*_ is the rate NICD enters the nucleus (production rate), Γ_*P*_ and Γ_*up*_ are the degradation rates of phosphorylated/unphosphorylated NICD, respectively, 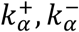 are the association and dissociation rates of the NCM complex to an SPS site, *k*_*p*_ is the Cdk8 phosphorylation rate of bound unphosphorylated NICD, and *N*_*ub*_ is the number of unbound SPS sites.

The total nuclear NICD concentration, *NICD*_*tot*_, is the sum of the phosphorylated, *NICD*_*p*_, and unphosphorylated fractions, *NICD*_*up*_:

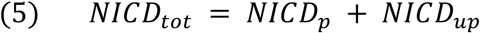

The total phosphorylated and unphosphorylated NICD are the sum of the bound (index *b*) and unbound (index *ub*) fractions:

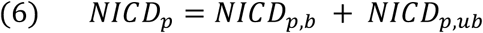

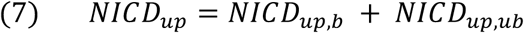

Combining equations 1-7 and assuming the system is in steady state gives:

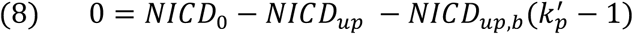

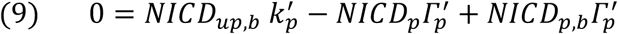

where we define the dimensionless parameters: 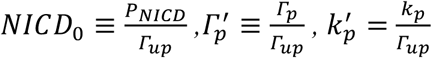.

*NICD*_*up,b*_ and *NICD*_*p,b*_ are calculated using a physical model based on equilibrium statistical mechanics (Brewster, et al., 2014). The conceptual basis of such models is that the occupancy of binding sites can be deduced by examining the equilibrium probabilities of binding and unbinding of transcription factors to transcription factor binding sites. In such models, each state of the system is denoted with a statistical weight (*S*_*i*_). In equilibrium, the statistical weights can be represented as the ratio of the concentration of each binding species [*X*], to the dissociation rate 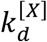 associated with that interaction so that 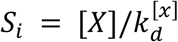 (White, et al., 2012). The partition function is defined as the summation of all possible statistical weights of the system:

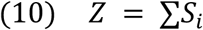

The partition function is the normalization factor by which the probabilities of the different states of the system are calculated so that the probability for state *j* is:

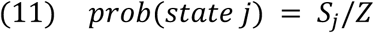

For our model, the statistical weights of the states of bound unphosphorylated and phosphorylated NICD are:

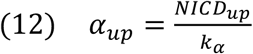

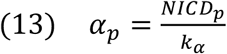

where 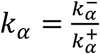 is the dissociation constant of NCM to an SPS site. Thus, the partition function of one binding site is:

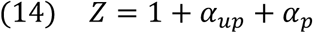

The total number of SPS sites, *N*, is comprised of endogenous SPS sites (*N*_*e*_) and synthetic SPS sites (*N*_*s*_):

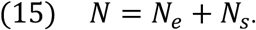

Note, *N*_*e*_ is an effective number for the cumulative impact of all endogenous sites (i.e. the many weak Notch binding sites within the genome) relative to the effect of the strong SPS sites in the *GBE-SPS* transgenes.

Combining equation 9-15 results in the following two equations:

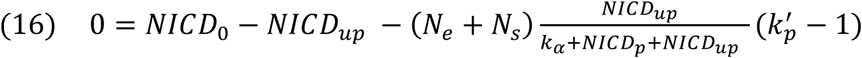

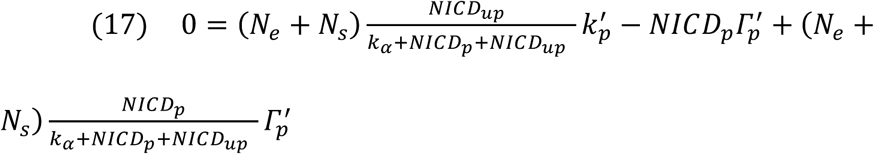

This statistical mechanics approach is based on three main assumptions: (1) The binding dynamics (on and off rates) are much faster than the dynamics determining the level of NICD in the nucleus. This is clearly valid as the DNA binding time scales are of the order of seconds (Gomez-Lamarca, et al., 2018) and degradation time scales are of the order of minutes to hours. (2) The number of NICD molecules in the nucleus is larger than the number of SPS sites. Since we typically look at a range of NICD concentrations of 10^2^ − 10^4^ per nucleus, and a maximum number of SPS sites of 36, this assumption is also justified. (3) For simplicity, we assume that binding to different SPS sites are independent.

We also consider the situation where the nuclear NICD concentration is much higher than the dissociation rate: [*NICD*_*p*_] + [*NICD*_*up*_] ≫ *k*_*α*_, namely, that we are in a strong binding regime. Under this assumption, the results are largely independent of the values of *k*_*α*_

We use equations (16) and (17) to solve for *NICD*_*p*_, *NICD*_*up*_, and *NICD*_*tot*_ and obtain their steady state levels for each set of parameters. These steady state solutions were used to plot Figures 4b and 4h (model curves).

#### Analysis of the linear regime

Since phosphorylation of NICD by Cdk8 occurs only for bound NICD, it can be assumed that for a low number of SPS sites *NICD*_*p*_, *k*_*α*_ (*N*_*e*_ + *N*_*s*_) ≪ *NICD*_*up*_. In this regime, equations 16 and 17 are approximated by:

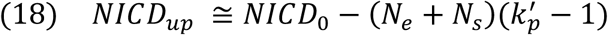

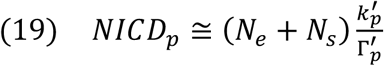

The total concentration of NICD is then:

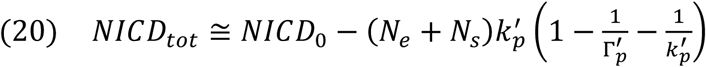

We now assume the phosphorylation rate of *NICD*_*up,b*_ is much faster than the degradation rate of *NICD*_*up*.*ub*_, that is: *k*_*p*_ ≫ Γ_*up*_. Under this assumption equation (20) becomes:

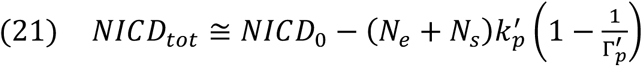

This analysis predicts that the slope in the linear regime is

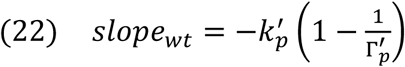

For mutant *skd* heterozygotes (*skd*^+/−^), we expect the phosphorylation rate to change to 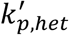. The expression for the total NICD in *skd*^+/−^ is then:

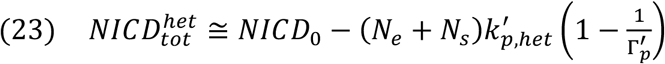

The ratio of the slopes between the wild type and *skd*^+/−^ will simply be:

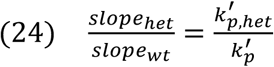

If *skd* is a limiting factor for the formation of the Cdk8 Mediator submodule, it is expected that reducing its copy number from 2 to 1 in *skd*^+/−^ would result in halving the Cdk8 phosphorylation activity, i.e. that 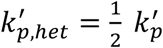 of the wild type.

The difference between equations 21 and 23 at *N*_*s*_ = 0 gives an expression for *N*_*e*_:

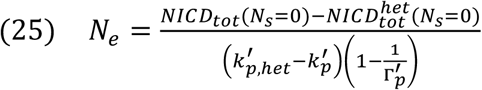

To check the ratio between wildtype and *skd* het slopes and to estimate *N*_*e*_, we performed linear regression on the data for the mean values of *6S-GFP* expression in Figure 4h using the first 3 points of wildtype data (the fourth point is in the saturated regime) and the 4 points of *skd*^+/−^ data. We note that the data is normalized to the mean fluorescence level of *6S-GFP* expression at *N*_*s*_ = 0, so equations 21 and 23 are normalized by *NICD*_*tot*_(*N*_*s*_ = 0). This normalization factor does not affect the expressions in equations 24 and 25 as it cancels out. The errors are estimated using standard error calculation on multivariate expression (Clifford, 1973).

#### Estimation of *k*_*p*_

The slope of the normalized linear fit is given by

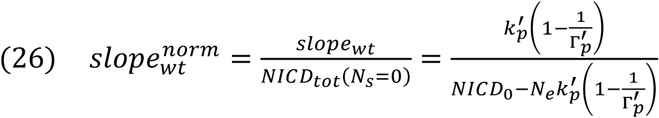

Which leads to the following expression for *k*_*p*_

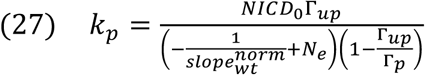

This expression allows estimating *k*_*p*_ for different parameter values. We use the calculated values of 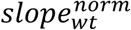 and *N*_*e*_. We estimate the steady state amount of NICD in the nucleus, *NICD*_0_, ranges between 100 (below that, the concentration is unlikely to activate multiple Notch targets in the nucleus) and 10,000. The upper limit is based on the fact that bicoid concentration is about 10,000 molecules/nucleus (Gregor, et al., 2007). Since endogenous NICD concentration is so small that it is notoriously hard to detect it in the nucleus using standard imaging techniques (Couturier, et al., 2012), we estimate that it is not larger than the typical concentration of Bicoid. The estimated range of unphosphorylated NICD is between 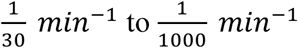 corresponding to half-lives of the range of 0.5-16 hours, which fits the typical half-lives of proteins. Note, that the analysis in Figure 3F shows a half-life of about 120 *min* in cell culture. Finally, since we assume that 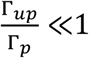 the exact value of Γ_*p*_ has only a weak effect on the values of *k*_*p*_. For the calculation we take it to be 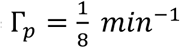 which is close to the rate observed in *Drosophila* cell culture (Housden, et al., 2013).

#### Dynamic simulations

To study the dynamics of NICD in the nucleus, we numerically solved the dynamic equations corresponding to equations 16 and 17:

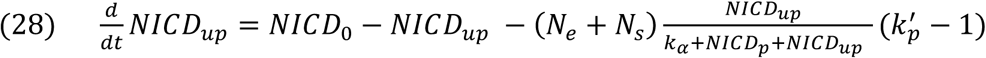

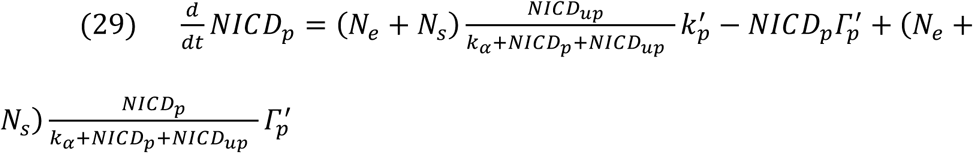

The equations were solved using ODE solver in MATLAB, with initial conditions *NICD*_*up*_(*t* = 0) = *NICD*_*p*_(*t* = 0) = 0. The values of parameters used for the simulations are given in Table S1. For simulating wildtype cells, we assumed *N*_*s*_ = 0 and *N*_*e*_ = 5.4. For simulating *N*^+/−^ cells, we assumed 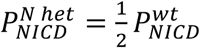. For simulating cells with two copies of 6SG, we assumed *N*_*s*_ = 12.

#### Parameter values

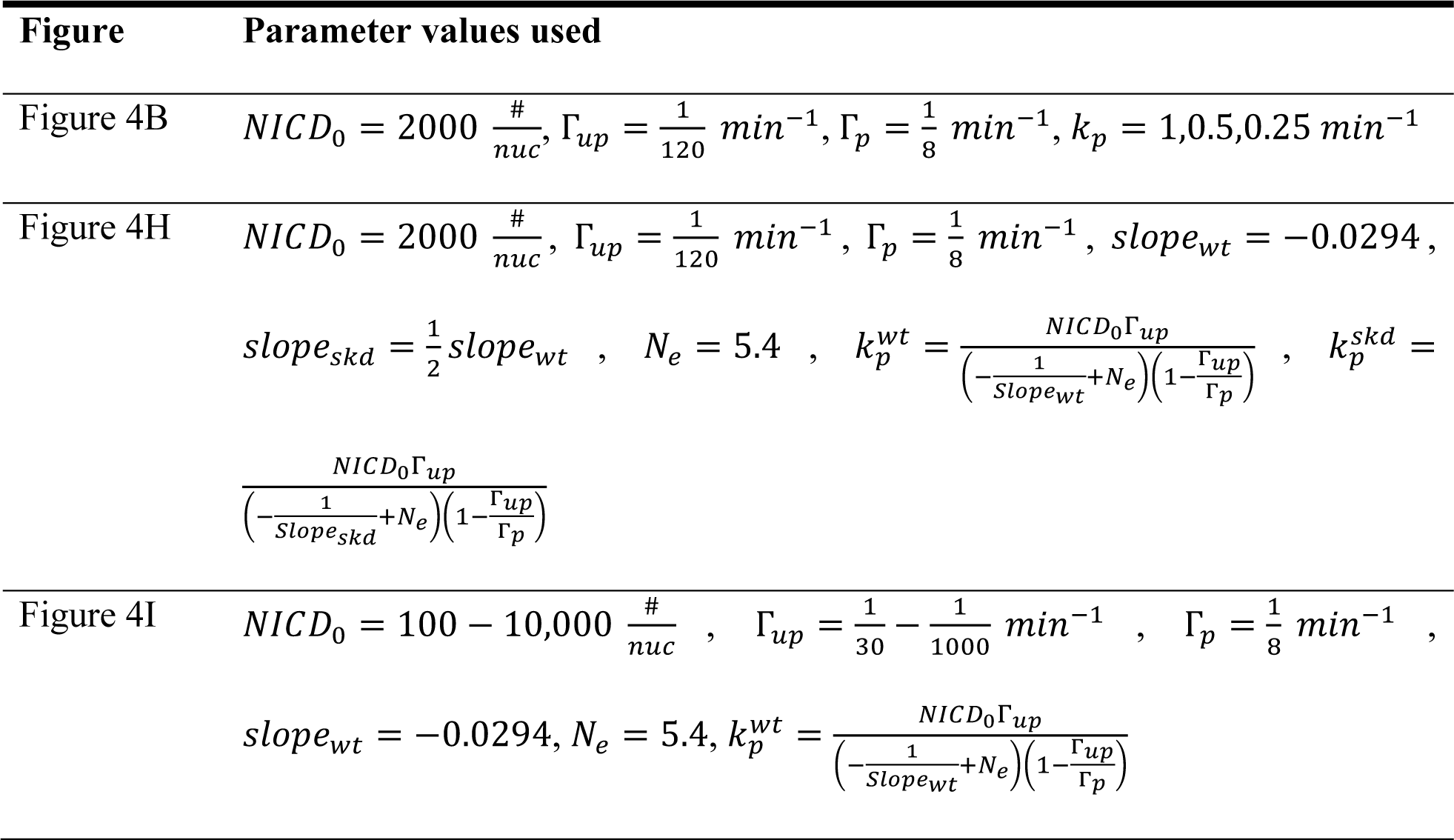
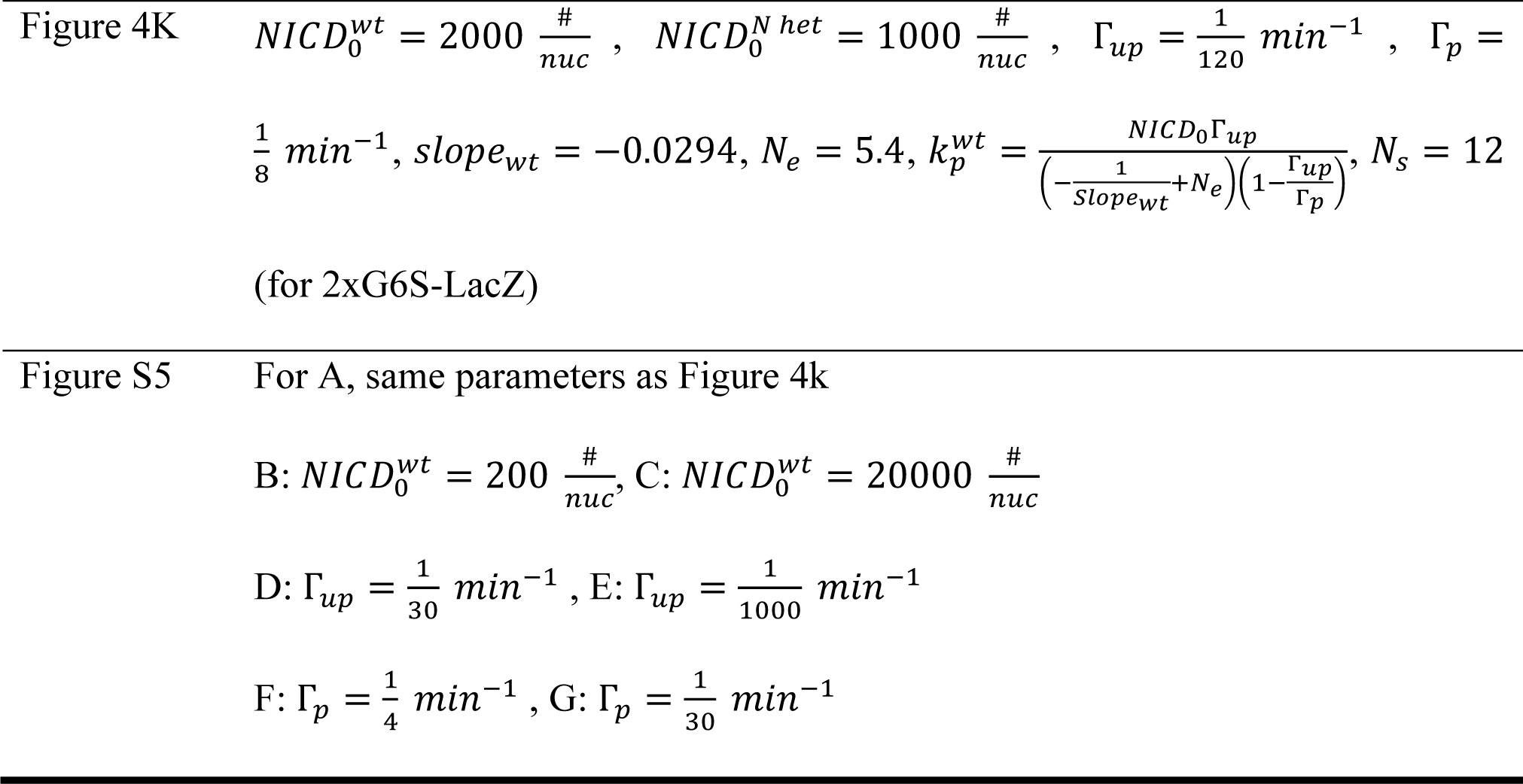

## LEAD CONTACT AND MATERIALS AVAILABILITY

All materials used in this study will be made freely available. Further information and requests for resources and reagents should be directed to and will be fulfilled by the Lead Contact, Brian Gebelein.

## DATA AND CODE AVAILABILITY

All simulation codes are available in the GitHub repository at https://github.com/OhadGolan/NICD-concentration-in-the-nucleus-as-by-binding-site-coupled-NICD-degradation.git.

