## Supplemental Information for "Enhancer architecture sensitizes cell specific responses to *Notch* gene dose via a bind and discard mechanism"

### SUPPLEMENTAL FIGURES

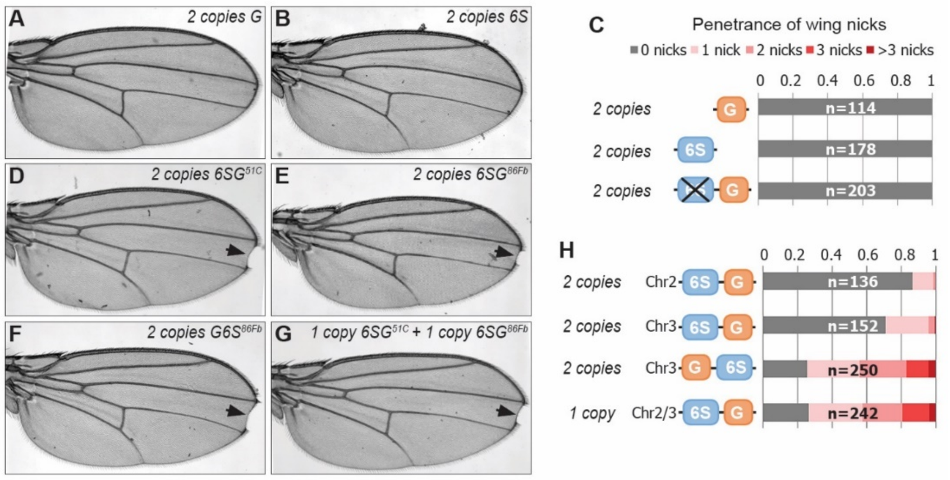


**Figure S1. Quantification GBE and SPS sites are both required to induce the formation of wing nicks. Related to Figure 1. A-B**. Wing images from flies containing either 2 copies of *3xGBE-lacZ* (A) or *6xSPS-lacZ* (B) reveals no wing notching phenotypes. **C**. Wing notching penetrance from flies with indicated genotypes. **D-G**. Wing images from flies with indicated genotypes. 51C and 86Fb are genomic loci on second and third chromosome, respectively and arrows denote wing notches. **H**. Wing notching penetrance and severity from flies with indicated genotypes. Number of wings scored (n) for each genotype is noted.

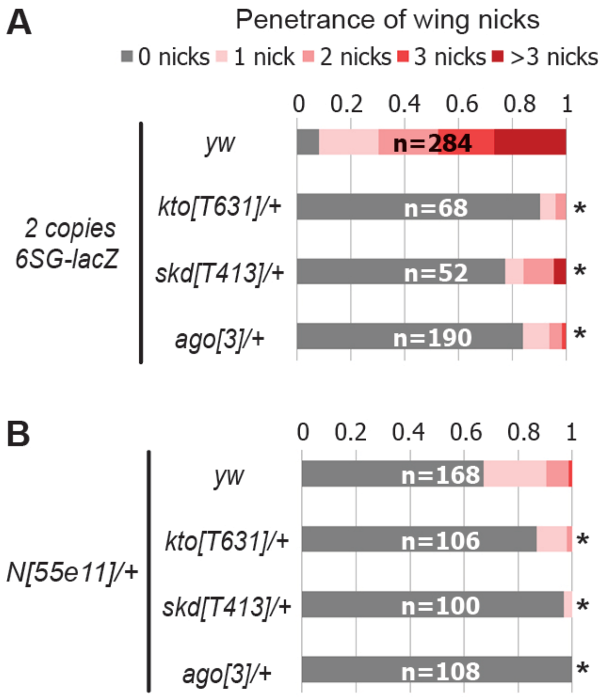


**Figure S2. Decreased gene dose of the Cdk8-Mediator submodule suppresses the formation of wing nicks. Related to Figure 3A-B. A**. Wing notching penetrance and severity from flies containing 2 copies of *6SG-lacZ* in wild-type (*yw*) or heterozygotes of components of the Cdk8-Mediator submodule or *Drosophila Fbw7* (*ago*). Wild-type data is the same as Figure 3a as it was performed in the same experiment (* $p < 0.05$). **B**. Wing notching penetrance and severity from *N* heterozygotes crossed to a wild-type (*yw*) or heterozygotes of either components of the Cdk8-Mediator submodule or *ago*. Wild-type data is the same as Figure 3b as it was performed in the same experiment. Number of wings scored (n) for each genotype is noted. Proportional odds model with Bonferroni adjustment (* $p < 0.05$).

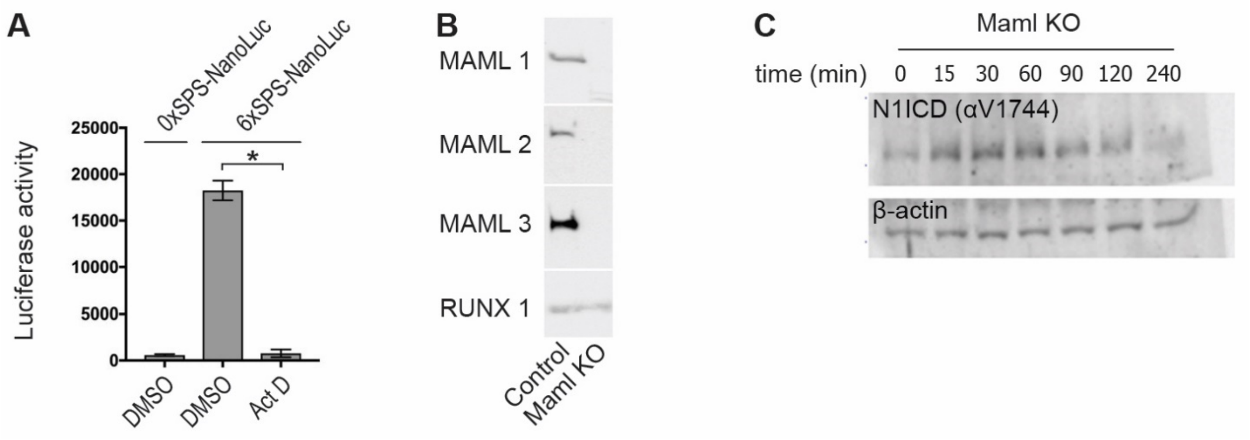


**Figure S3. Transcription inhibition doesn't impact NICD mobility. Related to Figure 3E-H. A**. Luciferase activity of *6xSPS-NanoLuc* (*6S*) after Notch activation in mK4 cells treated with either DMSO or Actinomycin D (50 nM). Empty luciferase vector (0xSPS-NanoLuc) was used as control (WT). One-way ANOVA analysis (* $p = 0.0002$). **B**. Western blot of MAML1-3 in either *wild-type* or *MAML*-deficient mK4 cells. RUNX1 is used as loading control. **C**. Western blot of N1ICD and β-actin after Notch activation in either *wild-type* or *MAML*-deficient mK4 cells. Note, N1ICD mobility from *MAML*-deficient cells did not change over time, consistent with post-translational modifications requiring MAML-mediated DNA binding.

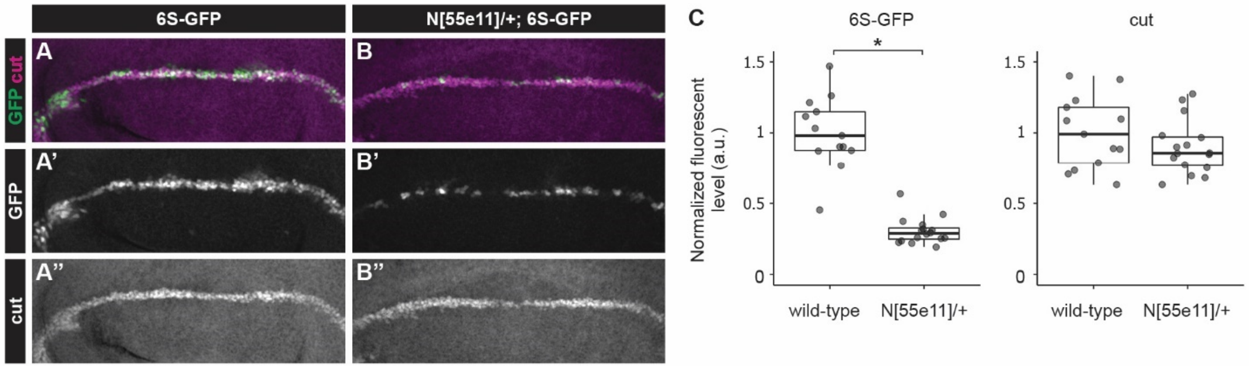


**Figure S4. *6S-GFP* reporter activity is sensitive to *Notch* gene dose. Related to Figure 4D-H. A-B”**. Representative imsmunostained third instar wing discs from *wild-type* or *N* heterozygous female flies containing *6xSPS-GFP* (*6S-GFP*) stained with cut (magenta). **C**. Quantification of GFP and Cut levels from wing discs of indicated genotype. Each dot represents the average wing margin cell pixel intensity as measured from an individual imaginal disc. Two-sided Student’s t-test (* $p < 0.05$).

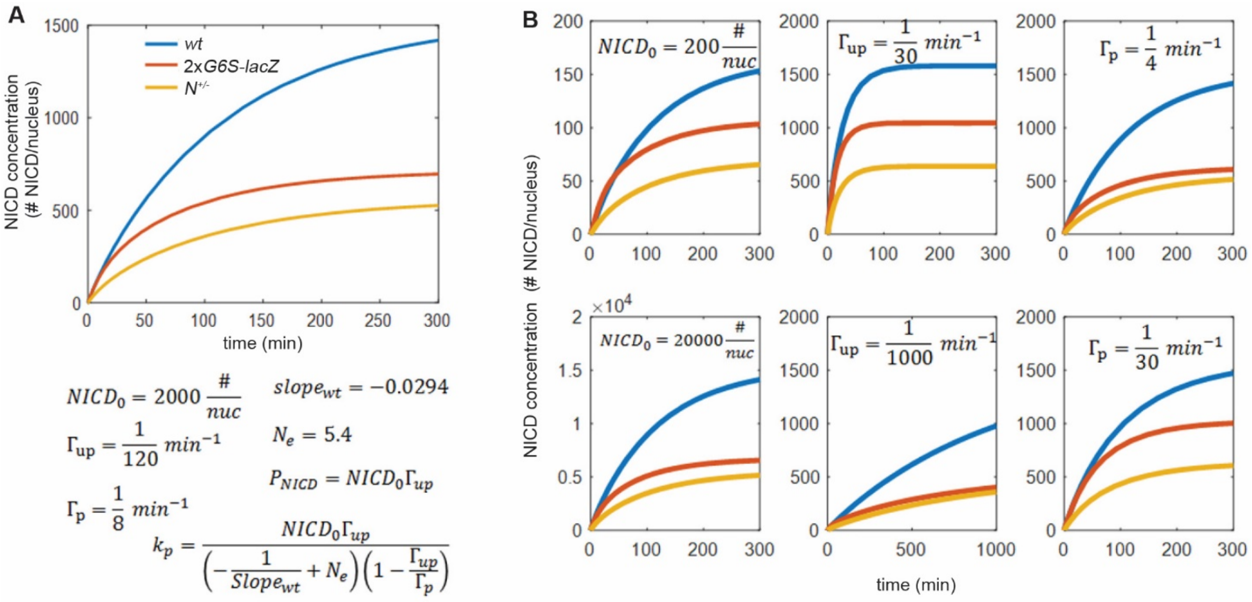


**Figure S5. Dynamic model is robust for to variations in parameters. Related to Figure 4K. A**. Top: Simulation of NICD levels as a function of time after onset of Notch activation (at t=0 min) in wild-type (blue), *N* heterozygotes (yellow) and *SPS-GBE* flies (red). Bottom: Parameters used for the model. The values of $slope_{wt}$ and $N_e$ were extracted from the analysis of Figure 4H. Other parameter values were chosen in the middle of the range estimation as described in the modeling section. **B**. Simulations of NICD levels as a function of time for different parameters as indicated. In each plot, one parameter is changed compared to (A) with all other parameters remaining the same. This analysis shows that varying $NICD_0$ and $\Gamma_{up}$ over two orders of magnitude, and $\Gamma_p$ over one order of magnitude do not affect the general conclusion that long duration processes, but not short duration processes, are sensitive to the CDK8-mediated degradation mechanism.

**Table S1. Genetic crosses performed to generate the analyzed progeny.**

| | Female | Male |
|---|---|---|
| Figure 1F | 6SG-lacZ$_{86Fb}$/6SG-lacZ$_{86Fb}$ | yw$^{1118}$ |
| Figure 1G | 6SG-lacZ$_{86Fb}$/6SG-lacZ$_{86Fb}$ | 6SG-lacZ$_{86Fb}$/6SG-lacZ$_{86Fb}$ |
| Figure 1H | 6SG-lacZ$_{51c}$/6SG-lacZ$_{51c}$;6SG-lacZ$_{86Fb}$/6SG-lacZ$_{86Fb}$ | 6SG-lacZ$_{51c}$/6SG-lacZ$_{51c}$;6SG-lacZ$_{86Fb}$/6SG-lacZ$_{86Fb}$ |
| Figure 1I | 12CG-lacZ$_{51c}$/12CG-lacZ$_{51c}$;12CG-lacZ$_{86Fb}$/12CG-lacZ$_{86Fb}$ | 12CG-lacZ$_{51c}$/12CG-lacZ$_{51c}$;12CG-lacZ$_{86Fb}$/12CG-lacZ$_{86Fb}$ |
| Figure 1J | G2S-lacZ$_{86Fb}$/G2S-lacZ$_{86Fb}$ | G2S-lacZ$_{86Fb}$/G2S-lacZ$_{86Fb}$ |
| | G6S-lacZ$_{86Fb}$/G6S-lacZ$_{86Fb}$ | G6S-lacZ$_{86Fb}$/G6S-lacZ$_{86Fb}$ |
| | G24S-lacZ$_{86Fb}$/G24S-lacZ$_{86Fb}$ | G24S-lacZ$_{86Fb}$/G24S-lacZ$_{86Fb}$ |
| Figure 1K | 6SG-lacZ$_{86Fb}$/6SG-lacZ$_{86Fb}$ | 6SG-lacZ$_{86Fb}$/6SG-lacZ$_{86Fb}$ |
| | 12CG-lacZ$_{86Fb}$/12CG-lacZ$_{86Fb}$ | 12CG-lacZ$_{86Fb}$/12CG-lacZ$_{86Fb}$ |
| Figure 2A | N$^{55e11}$ FRT19A/FM7c | yw$^{1118}$ |
| Figure 2B | N$^{55e11}$ FRT19A/FM7c | G6S-lacZ$_{86Fb}$/G6S-lacZ$_{86Fb}$ |
| Figure 2C | N$^{55e11}$ FRT19A/FM7c | yw$^{1118}$ |
| | N$^{55e11}$ FRT19A/FM7c | 6SG-lacZ$_{86Fb}$/6SG-lacZ$_{86Fb}$ |
| | yw$^{1118}$ | 6SG-lacZ$_{86Fb}$/6SG-lacZ$_{86Fb}$ |
| Figure 2D | PBac{N-GFP.FLAG}/PBac{N-GFP.FLAG} | PBac{N-GFP.FLAG}/PBac{N-GFP.FLAG} |
| Figure 2E | 6SG-lacZ$_{51c}$/6SG-lacZ$_{51c}$;PBac{N-GFP.FLAG}/PBac{N-GFP.FLAG} | 6SG-lacZ$_{51c}$/6SG-lacZ$_{51c}$;PBac{N-GFP.FLAG}/PBac{N-GFP.FLAG} |
| Figure 2F | 6SG-lacZ$_{51c}$/6SG-lacZ$_{51c}$ | 6SG-lacZ$_{51c}$/6SG-lacZ$_{51c}$ |
| | 6SG-lacZ$_{51c}$/6SG-lacZ$_{51c}$;PBac{N-GFP.FLAG}/PBac{N-GFP.FLAG} | 6SG-lacZ$_{51c}$/6SG-lacZ$_{51c}$;PBac{N-GFP.FLAG}/PBac{N-GFP.FLAG} |
| | 6SG-lacZ$_{51c}$/6SG-lacZ$_{51c}$;6SG-lacZ$_{86Fb}$/6SG-lacZ$_{86Fb}$ | H$^{1}$/TM6B, Tb$^{1}$ |
| Figure 2G | yw$^{1118}$ | H$^{1}$/TM6B, Tb$^{1}$ |

| | | |
|---|---|---|
| Figure 2H | 6SG-lacZ$_{51c}$/6SG-lacZ$_{51c}$;6SG-lacZ$_{86Fb}$/6SG-lacZ$_{86Fb}$ | $H^{1}$/TM6B, $Tb^{1}$ |
| Figure 2I | $yw^{1118}$ | $H^{1}$/TM6B, $Tb^{1}$ |
| | 6SG-lacZ$_{51c}$/6SG-lacZ$_{51c}$;6SG-lacZ$_{86Fb}$/6SG-lacZ$_{86Fb}$ | $H^{1}$/TM6B, $Tb^{1}$ |
| | 12CG-lacZ$_{51c}$/12CG-lacZ$_{51c}$;12CG-lacZ$_{86Fb}$/12CG-lacZ$_{86Fb}$ | $H^{1}$/TM6B, $Tb^{1}$ |
| Figure 2J | $N^{55e11}$ FRT19A/FM7c | $yw^{1118}$ |
| Figure 2K | G6S-lacZ$_{86Fb}$/G6S-lacZ$_{86Fb}$ | G6S-lacZ$_{86Fb}$/G6S-lacZ$_{86Fb}$ |
| Figure 2L | $N^{55e11}$ FRT19A/FM7c | $yw^{1118}$ |
| | $N^{55e11}$ FRT19A/FM7c | G6S-lacZ$_{86Fb}$/G6S-lacZ$_{86Fb}$ |
| | 6SG-lacZ$_{51c}$/6SG-lacZ$_{51c}$;6SG-lacZ$_{86Fb}$/6SG-lacZ$_{86Fb}$ | $yw^{1118}$ |
| | $yw^{1118}$ | $H^{1}$/TM6B, $Tb^{1}$ |
| | 6SG-lacZ$_{51c}$/6SG-lacZ$_{51c}$;6SG-lacZ$_{86Fb}$/6SG-lacZ$_{86Fb}$ | $H^{1}$/TM6B, $Tb^{1}$ |
| Figure 3A | 6SG-lacZ$_{51c}$/6SG-lacZ$_{51c}$;6SG-lacZ$_{86Fb}$/6SG-lacZ$_{86Fb}$ | $yw^{1118}$ |
| | 6SG-lacZ$_{51c}$/6SG-lacZ$_{51c}$;6SG-lacZ$_{86Fb}$/6SG-lacZ$_{86Fb}$ | FRT80 $cdk8^{K185}$/TM6B |
| | 6SG-lacZ$_{51c}$/6SG-lacZ$_{51c}$;6SG-lacZ$_{86Fb}$/6SG-lacZ$_{86Fb}$ | FRT82 $cycC^{Y5}$/TM6B |
| | 6SG-lacZ$_{51c}$/6SG-lacZ$_{51c}$;6SG-lacZ$_{86Fb}$/6SG-lacZ$_{86Fb}$ | $kto^{T241}$ FRT80B/TM6B, $Tb^{1}$ |
| | 6SG-lacZ$_{51c}$/6SG-lacZ$_{51c}$;6SG-lacZ$_{86Fb}$/6SG-lacZ$_{86Fb}$ | $skd^{T13}$ FRT80B/TM6B, $Tb^{1}$ |
| | 6SG-lacZ$_{51c}$/6SG-lacZ$_{51c}$;6SG-lacZ$_{86Fb}$/6SG-lacZ$_{86Fb}$ | $ago^{1}$ FRT80B/TM6B, $Tb^{1}$ |
| Figure 3B | $N^{55e11}$ FRT19A/FM7c | $yw^{1118}$ |
| | $N^{55e11}$ FRT19A/FM7c | FRT80 $cdk8^{K185}$/TM6B |

| | | |
|---|---|---|
| | *$N^{55e11}$ FRT19A/FM7c* | *FRT82 $cycC^{Y5}$/TM6B* |
| | *$N^{55e11}$ FRT19A/FM7c* | *$kto^{T241}$ FRT80B/TM6B, $Tb^{1}$* |
| | *$N^{55e11}$ FRT19A/FM7c* | *$skd^{T13}$ FRT80B/TM6B, $Tb^{1}$* |
| | *$N^{55e11}$ FRT19A/FM7c* | *$ago^{1}$ FRT80B/TM6B, $Tb^{1}$* |
| Figure 3D | *G24S-$GFP_{51c}$/G24S-$GFP_{51c}$;G24S-$GFP_{86Fb}$/G24S-$GFP_{86Fb}$* | *$yw^{1118}$* |
| | *$G24S_{51c}$/$G24S_{51c}$; $G24S_{86Fb}$/$G24S_{86Fb}$* | *$yw^{1118}$* |
| | *G24S-$GFP_{51c}$/G24S-$GFP_{51c}$;G24S-$GFP_{86Fb}$/G24S-$GFP_{86Fb}$* | *$skd^{T413}$ FRT80B/TM6B, $Tb^{1}$* |
| | *$G24S_{51c}$/$G24S_{51c}$; $G24S_{86Fb}$/$G24S_{86Fb}$* | *$skd^{T413}$ FRT80B/TM6B, $Tb^{1}$* |
| Figure 4C | *6S-$GFP_{22A}$/6S-$GFP_{22A}$;G6S-$lacZ_{86Fb}$/G6S-$lacZ_{86Fb}$* | *$yw^{1118}$* |
| | *6S-$GFP_{22A}$/6S-$GFP_{22A}$;G6S-$lacZ_{86Fb}$/G6S-$lacZ_{86Fb}$* | *$skd^{T413}$ FRT80B/TM6B, $Tb^{1}$* |
| | *6S-$GFP_{22A}$/6S-$GFP_{22A}$;(G6S)2-$lacZ_{86Fb}$/(G6S)2-$lacZ_{86Fb}$* | *$yw^{1118}$* |
| | *6S-$GFP_{22A}$/6S-$GFP_{22A}$;(G6S)2-$lacZ_{86Fb}$/(G6S)2-$lacZ_{86Fb}$* | *$skd^{T413}$ FRT80B/TM6B, $Tb^{1}$* |
| | *6S-$GFP_{22A}$/6S-$GFP_{22A}$;(G6S)3-$lacZ_{86Fb}$/(G6S)3-$lacZ_{86Fb}$* | *$yw^{1118}$* |
| | *6S-$GFP_{22A}$/6S-$GFP_{22A}$;(G6S)3-$lacZ_{86Fb}$/(G6S)3-$lacZ_{86Fb}$* | *$skd^{T413}$ FRT80B/TM6B, $Tb^{1}$* |
| Figure 4D | *6SG-$lacZ_{51c}$/6SG-$lacZ_{51c}$;G-$lacZ_{86Fb}$/G-$lacZ_{86Fb}$* | *$yw^{1118}$* |
| Figure 4E | *6S-$GFP_{22A}$/6S-$GFP_{22A}$;G6S-$lacZ_{86Fb}$/G6S-$lacZ_{86Fb}$* | *$yw^{1118}$* |
| Figure 4F | *6S-$GFP_{22A}$/6S-$GFP_{22A}$;(G6S)2-$lacZ_{86Fb}$/(G6S)2-$lacZ_{86Fb}$* | *$yw^{1118}$* |
| Figure 4G | *6S-$GFP_{22A}$/6S-$GFP_{22A}$;(G6S)3-$lacZ_{86Fb}$/(G6S)3-$lacZ_{86Fb}$* | *$yw^{1118}$* |
| Figure 4J | *$N^{55e11}$ FRT19A/FM7c* | *$yw^{1118}$* |
| | *$N^{55e11}$ FRT19A/FM7c* | *FRT80 $cdk8^{K185}$/TM6B* |

| | | |
|---|---|---|
| | *N*$^{55e11}$ *FRT19A/FM7c* | *FRT82 cycC*$^{Y5}$*/TM6B* |
| | *N*$^{55e11}$ *FRT19A/FM7c* | *kto*$^{T241}$ *FRT80B/TM6B, Tb*$^{1}$ |
| | *N*$^{55e11}$ *FRT19A/FM7c* | *skd*$^{T13}$ *FRT80B/TM6B, Tb*$^{1}$ |
| | *H*$^{1}$*/TM6B, Tb*$^{1}$ | *yw*$^{1118}$ |
| | *H*$^{1}$*/TM6B, Tb*$^{1}$ | *FRT80 cdk8*$^{K185}$*/TM6B* |
| | *H*$^{1}$*/TM6B, Tb*$^{1}$ | *FRT82 cycC*$^{Y5}$*/TM6B* |
| | *H*$^{1}$*/TM6B, Tb*$^{1}$ | *kto*$^{T241}$ *FRT80B/TM6B, Tb*$^{1}$ |
| | *H*$^{1}$*/TM6B, Tb*$^{1}$ | *skd*$^{T13}$ *FRT80B/TM6B, Tb*$^{1}$ |
| Figure S1A | *G-lacZ*$_{86Fb}$*/G-lacZ*$_{86Fb}$ | *G-lacZ*$_{86Fb}$*/G-lacZ*$_{86Fb}$ |
| Figure S1B | *6S-lacZ*$_{86Fb}$*/6S-lacZ*$_{86Fb}$ | *6S-lacZ*$_{86Fb}$*/6S-lacZ*$_{86Fb}$ |
| Figure S1C | *G-lacZ*$_{86Fb}$*/G-lacZ*$_{86Fb}$ | *G-lacZ*$_{86Fb}$*/G-lacZ*$_{86Fb}$ |
| | *6S-lacZ*$_{86Fb}$*/6S-lacZ*$_{86Fb}$ | *6S-lacZ*$_{86Fb}$*/6S-lacZ*$_{86Fb}$ |
| | *6SmutG-lacZ*$_{86Fb}$*/6SmutG-lacZ*$_{86Fb}$ | *6SmutG-lacZ*$_{86Fb}$*/6SmutG-lacZ*$_{86Fb}$ |
| Figure S1D | *6SG-lacZ*$_{51c}$*/6SG-lacZ*$_{51c}$ | *6SG-lacZ*$_{51c}$*/6SG-lacZ*$_{51c}$ |
| Figure S1E | *6SG-lacZ*$_{86Fb}$*/6SG-lacZ*$_{86Fb}$ | *6SG-lacZ*$_{86Fb}$*/6SG-lacZ*$_{86Fb}$ |
| Figure S1F | *G6S-lacZ*$_{86Fb}$*/G6S-lacZ*$_{86Fb}$ | *G6S-lacZ*$_{86Fb}$*/G6S-lacZ*$_{86Fb}$ |
| Figure S1G | *6SG-lacZ*$_{51c}$*/6SG-lacZ*$_{51c}$*;6SG-lacZ*$_{86Fb}$*/6SG-lacZ*$_{86Fb}$ | *yw*$^{1118}$ |
| Figure S1H | *6SG-lacZ*$_{51c}$*/6SG-lacZ*$_{51c}$ | *6SG-lacZ*$_{51c}$*/6SG-lacZ*$_{51c}$ |
| | *6SG-lacZ*$_{86Fb}$*/6SG-lacZ*$_{86Fb}$ | *6SG-lacZ*$_{86Fb}$*/6SG-lacZ*$_{86Fb}$ |
| | *G6S-lacZ*$_{86Fb}$*/G6S-lacZ*$_{86Fb}$ | *G6S-lacZ*$_{86Fb}$*/G6S-lacZ*$_{86Fb}$ |
| | *6SG-lacZ*$_{51c}$*/6SG-lacZ*$_{51c}$*;6SG-lacZ*$_{86Fb}$*/6SG-lacZ*$_{86Fb}$ | *yw*$^{1118}$ |
| Figure S2A | *6SG-lacZ*$_{51c}$*/6SG-lacZ*$_{51c}$*;6SG-lacZ*$_{86Fb}$*/6SG-lacZ*$_{86Fb}$ | *yw*$^{1118}$ |

| | | |
|---|---|---|
| | *6SG-lacZ*$_{51c}$*/6SG-lacZ*$_{51c}$*;6SG-lacZ*$_{86Fb}$*/6SG-lacZ*$_{86Fb}$ | *kto*$^{T631}$ *FRT80B/TM6B, Tb*$^{1}$ |
| | *6SG-lacZ*$_{51c}$*/6SG-lacZ*$_{51c}$*;6SG-lacZ*$_{86Fb}$*/6SG-lacZ*$_{86Fb}$ | *skd*$^{T413}$ *FRT80B/TM6B, Tb*$^{1}$ |
| | *6SG-lacZ*$_{51c}$*/6SG-lacZ*$_{51c}$*;6SG-lacZ*$_{86Fb}$*/6SG-lacZ*$_{86Fb}$ | *ago*$^{3}$ *FRT80B/TM6B, Tb*$^{1}$ |
| Figure S2B | *N*$^{55e11}$ *FRT19A/FM7c* | *yw*$^{1118}$ |
| | *N*$^{55e11}$ *FRT19A/FM7c* | *kto*$^{T631}$ *FRT80B/TM6B, Tb*$^{1}$ |
| | *N*$^{55e11}$ *FRT19A/FM7c* | *skd*$^{T413}$ *FRT80B/TM6B, Tb*$^{1}$ |
| | *N*$^{55e11}$ *FRT19A/FM7c* | *ago*$^{3}$ *FRT80B/TM6B, Tb*$^{1}$ |
| Figure S4A | *yw*$^{1118}$ | *6S-GFP*$_{86Fb}$*/6S-GFP*$_{86Fb}$ |
| | *N*$^{55e11}$ *FRT19A/FM7i, ActGFP* | *6S-GFP*$_{86Fb}$*/6S-GFP*$_{86Fb}$ |
